# Discovering and exploiting multiple types of DNA methylation from individual bacteria and microbiome using nanopore sequencing

**DOI:** 10.1101/2020.02.18.954636

**Authors:** Alan Tourancheau, Edward A. Mead, Xue-Song Zhang, Gang Fang

## Abstract

Nanopore sequencing provides a great opportunity for direct detection of chemical DNA modification. However, existing computational methods were either trained for detecting a specific form of DNA modification from one, or a few, specific sequence contexts (*e.g.* 5-methylcytosine from CpG dinucleotides) or for allowing *de novo* detection without effectively differentiating between different forms of DNA modifications. As a result, none of these methods supports *de novo*, systematic study of unknown bacterial methylomes. In this work, by examining three types of DNA methylation in a large diversity of sequence contexts, we observed that nanopore sequencing signal displays complex heterogeneity across methylation events of the same type. To capture this complexity and enable nanopore sequencing for broadly applicable methylation discovery, we generated a training dataset from an assortment of bacterial species and developed a novel method that couples the identification and fine mapping of the three forms of DNA methylation into a multi-label classification design. We evaluated the method and then applied it to individual bacteria and mouse gut microbiome for reliable methylation discovery. In addition, we demonstrated in the microbiome analysis the use of DNA methylation for binning metagenomic contigs, associating mobile genetic elements with their host genomes, and for the first time, identifying misassembled metagenomic contigs. This novel method has broad utility for discovering different forms of DNA methylation from bacteria, assisting functional studies of epigenetic regulation in bacteria, and exploiting bacterial epigenomes for more effective metagenomic analyses.

## Introduction

Third generation sequencing, represented by Single Molecule Real-Time (SMRT) and nanopore sequencing, provides a great opportunity for the direct detection of multiple types of DNA methylation and other chemical modifications to DNA^1^. In SMRT sequencing, the instrument monitors not only the pulse fluorescence associated with each nucleotide, but also records the time it takes for the DNA polymerase to translocate from one nucleotide to the next, termed inter-pulse duration (IPD). Deviation of IPD, calculated by comparing native DNA with methylation free DNA (*e.g.* produced by whole genome amplification, WGA), is correlated with the presence of some DNA modifications^2^. In nanopore sequencing, chemical modification to DNA in the native library can affect current signals near modified bases, which can be detected using a similar native versus WGA design^3, 4^. While SMRT sequencing has already played a foundational role in the recent bloom of bacterial methylome studies^1, 5^ as well as in the study of eukaryotic methylomes^6–8^, the application of nanopore sequencing in mapping DNA methylation events are under active development^1^.

Great progress has been made in methods development for DNA modification detection using nanopore sequencing. Two early studies showed differences in current at multiple consecutive positions near the modified base when comparing nanopore sequencing signals from the same genomic regions with or without DNA methylation^3, 4^. Recently, a method using hidden Markov model (HMM) trained to detect 5- methylcytosine (5mC) in CpG islands was successfully applied to 5mC calling from human genome^9^. Another method coupling a hierarchical Dirichlet process with HMM demonstrated improved performance for detecting 5mC, 5-hydroxymethylcytosine (5hmC) and N6-methyladenine (6mA) on synthetic oligonucleotides or specific methylation motifs in the *E. coli* genome (C5mCWGG, and G6mATC)^10^. Three recent works developed classification models that can capture the difference between methylated sites (6mA/5mC) and non-methylated A’s/C’s^,11-13^. While encouraging, these model-based approaches remain limited in their ability to *de novo* identify diverse modification types at various sequence motifs. Following the design first proposed for SMRT sequencing^2^, two studies described nanopore-based methylation detection methods using a statistical comparison of current signal between native and methylation-free WGA DNA samples^14, 15^. While these methods have the advantage of not requiring *a priori* knowledge of the impact of specific types of DNA modification on ionic current, they do not *de novo* identify the specific methylation type or the precise modified position^16^.

To summarize, existing methods were either trained for detecting a specific type of DNA methylation from one of few specific sequence contexts (*e.g.* 5mC at CpG or 6mA at GATC) or allow more general detection without effectively differentiating between different forms of DNA methylation or identifying the exact modified position. To date, none of these methods have been applied to *de novo* characterize unknown bacterial methylomes in full extent (*i.e. de novo* methylation detection of all three major types of DNA methylation in bacteria), *de novo* identification of methylation type (*i.e.* assigning methylation type: 4mC, 5mC or 6mA), and *de novo* fine mapping of the methylated nucleotide.

In this work, by examining three types of DNA methylation (4mC, 5mC, and 6mA) in a large diversity of sequence context, we observed large variation and complex heterogeneity in terms of their impact on ionic current levels captured in nanopore sequencing. This observation has important implications suggesting that detection methods are best developed using a diverse collection of species. Bacterial epigenomes are highly motif-driven given the nature of restriction modification systems; nearly all occurrences (mostly >95%; often >99%) of a motif recognized by an active methyltransferase in a given prokaryotic genome are methylated^1, 17, 18^. Following this rationale, we built a training dataset and developed a novel, extensible method for *de novo* methylation typing and fine mapping of the three forms of DNA methylation at constitutively methylated motifs using a multi-label classification framework. We evaluated the methods and then applied it to individual bacteria and mouse gut microbiomes for reliable methylation discovery. For microbiome analysis, we also demonstrated the use of DNA methylation information for binning metagenomic contigs, associating mobile genetic elements with their host genomes, and for the first time, identifying misassembled metagenomic contigs.

## Results

### Heterogeneous signal variation induced by DNA methylation in nanopore sequencing

In the bacterial kingdom, DNA methylation has three primary forms: 6mA, 4mC and 5mC, all of which occur in a highly motif-driven manner: on average, each bacterial genome contains three methylation motifs, and nearly every occurrence of the target motifs is methylated^1, 5^. In order to comprehensively examine the variation of different types of DNA methylation within a broad scope of sequence context as measured by nanopore sequencing, we collected 46 well-characterized unique methylation motifs from a set of bacterial species with diverse methylation motifs (**Supplementary Table 1**; Methods). According to a previous study^5^ and REBASE curated database^19^ (Methods), these strains have a total of 46 unique and confident methylation motifs covering the three major methylation types (6mA motifs: 28; 4mC motifs: 7; 5mC motifs: 11; 308,773 methylation sites in total; **Fig. 1**; **Supplementary Table 2**). Nanopore sequencing was conducted on MinION with R9.4 flow cells achieving 175x coverage on average (**Supplementary Table 3**) for both the native DNA samples and their WGA samples.

**Figure 1:**
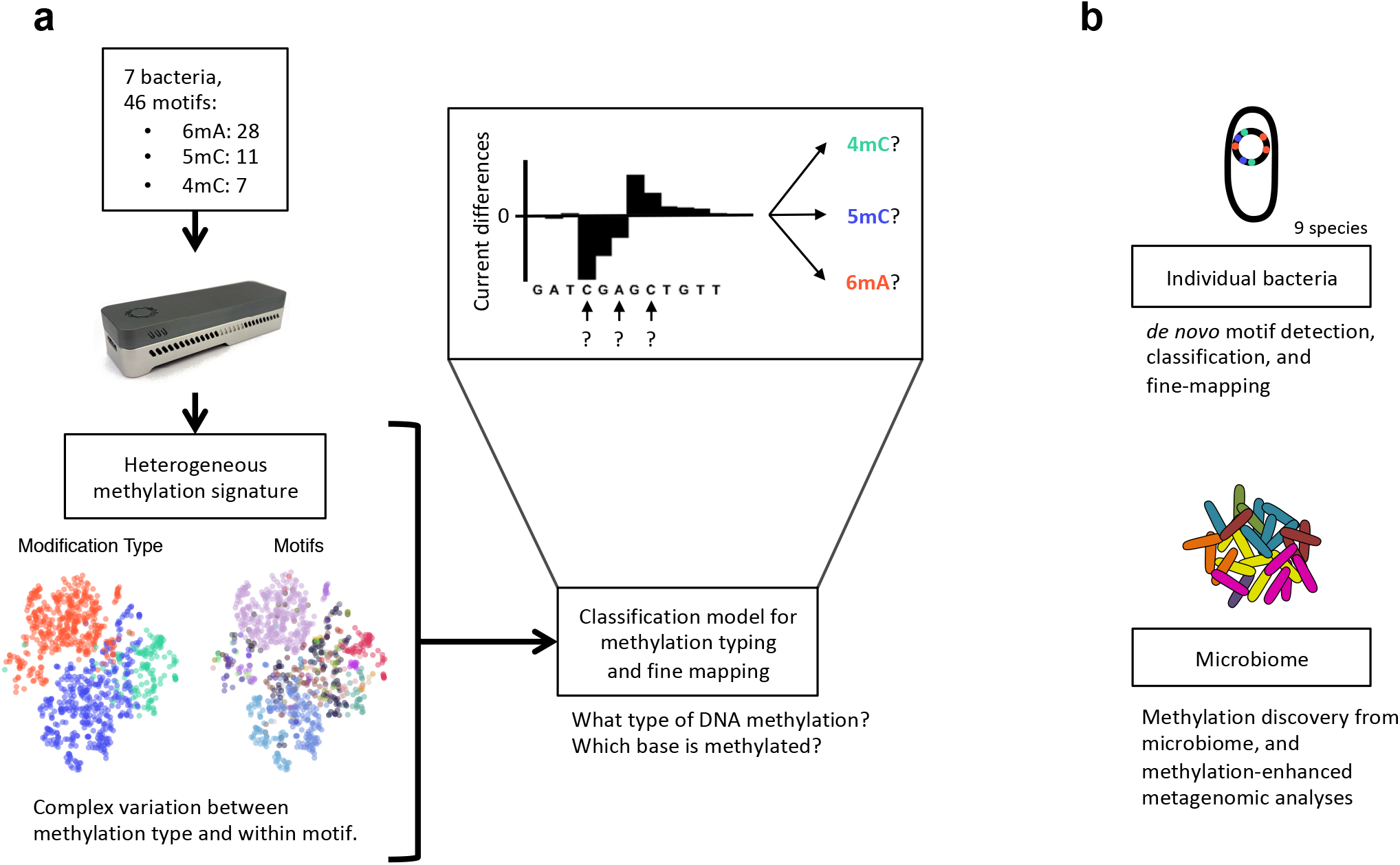
Schematics for method design and applications. (**a**) Using isolated bacteria with a wide variety of methylation motifs we explore the signal of DNA methylation in nanopore sequencing and characterize the major types of DNA methylation (4mC, 5mC, and 6mA). We observed a large variation and complex heterogeneity of current differences (native versus WGA) between methylation sequence context, which motivated us to develop a broadly applicable method for classifying DNA methylation into specific methylation type (4mC, 5mC, and 6mA) and fine mapping of the methylated base. (**b**) We performed comprehensive method evaluation and demonstrated the application of our method for methylation discovery from individual bacterial species (7 + 2 species) and microbiomes (methylation motif detection, classification, and fine mapping), as well as methylation-assisted metagenomic analysis (methylation binning and misassembly identification).

Read events and associated current levels (picoampere, pA) were aligned to reference genomes using Nanopolish^9^. After normalization and filtering, current differences between native and WGA datasets were computed for each genomic position (Methods). To examine the variation of current differences across different DNA methylation types and motifs, we extracted current differences around each methylated base ([-6 bp, +7 bp]) and computed the methylation motif signatures (distribution of current differences at relative distance from methylation motif, see Methods; **Fig. 2a**). Generally, the widths and amplitudes of perturbation in the methylation motif signatures vary between different motifs and methylation types (**Supplementary Fig. 1a-c**). The broadness of signal perturbation suggests that methylation induces current differences across multiple flanking bases, essentially due to DNA methylation disturbing the ionic current of multiple consecutive events while ratcheting through the nanopore^3, 4^. It is worth noting that this broadness contrasts with the deviations of kinetic DNA polymerase confined to a single base for 4mC and 6mA in SMRT sequencing^2, 20–23^.

**Figure 2:**
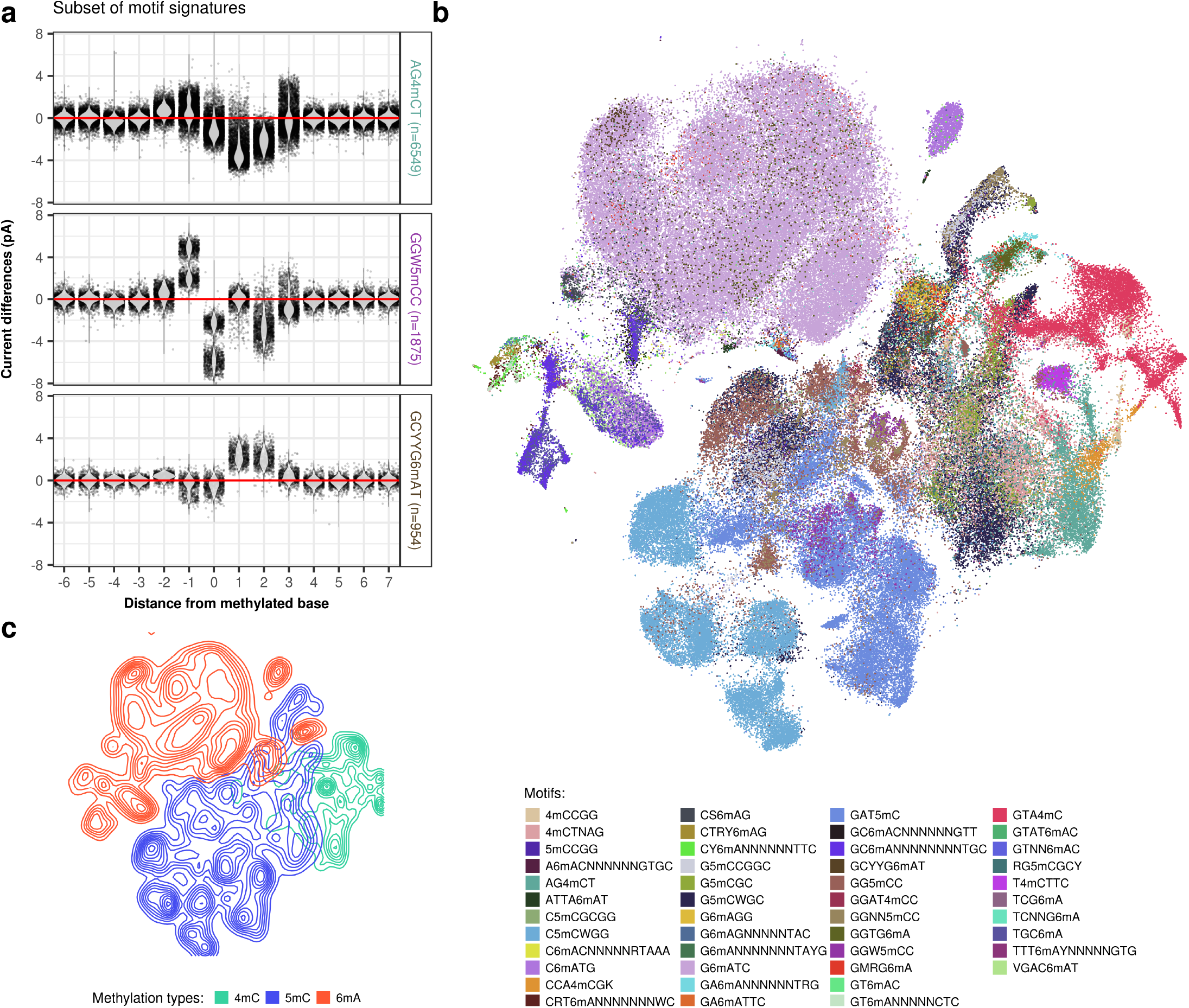
Systematic examination of three main types of DNA methylation with nanopore sequencing. (**a**) Variation of current differences across methylation occurrences as illustrated by motif signatures from three motifs (AG4mCT, GGW5mCC, and GCYYG6mAT). For each motif, current differences near methylated bases ([- 6 bp, + 7 bp]) from all isolated occurrences are plotted with conservation of relative distances to methylated bases. Distributions of current differences for each relative distance are displayed as a violin plot. Current differences axis is limited to −8 to 8 pA range. (**b**) Variation of current differences across methylation occurrences as illustrated by projection with t-SNE for 46 well-characterized motifs (**Supplementary Table 2**). Each dot represents one isolated motif occurrence colored by methylation motif. For each motif occurrence, current differences from 22 positions near methylated bases ([- 10 bp, + 11 bp]) were used. (**c**) Similar to **b** but colored by DNA methylation type with additional processing to reveal cluster density indicated by relief.

To obtain an overall view of the current differences across all the methylation types and methylation motifs, we subjected the 14 bp vectors ([-6 bp, +7 bp]) capturing current differences across 183,818 non-overlapping methylation motif occurrences to t-distributed stochastic neighbor embedding (t-SNE)^24^, a nonlinear dimensionality reduction algorithm (**Fig. 2b,c**, **Supplementary Fig. 2**). There is a general clustering pattern where methylation motif occurrences from the same methylation type tend to cluster together (**Fig. 2c** and **Supplementary Fig. 2b**), although there are apparent overlaps. Importantly, we observed that current differences associated with different methylation motifs of the same methylation type often form different clusters, some individual motifs form distinct sub-clusters, *i.e.* current differences generally varies between different motifs of the same methylation type (e.g. T4mCTTC and GTA4mC; **Fig. 2c** and **Supplementary Fig. 2b**), and even between methylation events within the same methylation motif (e.g. GGW5mCC; **Fig. 2a,b** and **Supplementary Fig. 2a**). Further analysis of signatures for subsets of the same motif suggests that this across-motif and within-motif variation can be in a large part explained by sequence variation from degenerated position in motifs as well as sequences flanking the consensus motifs. In **Fig. 3a,b**, we showed an illustrative example where signature sub-clusters for a 5mC motif (GGW5mCC) can be partially explained by sequence diversity near methylated bases (within-motif sequence variation). Similar observations were made with respect to sequence variation outside of consensus methylation motif (e.g. GAT5mC; **Fig. 3c**).

**Figure 3:**
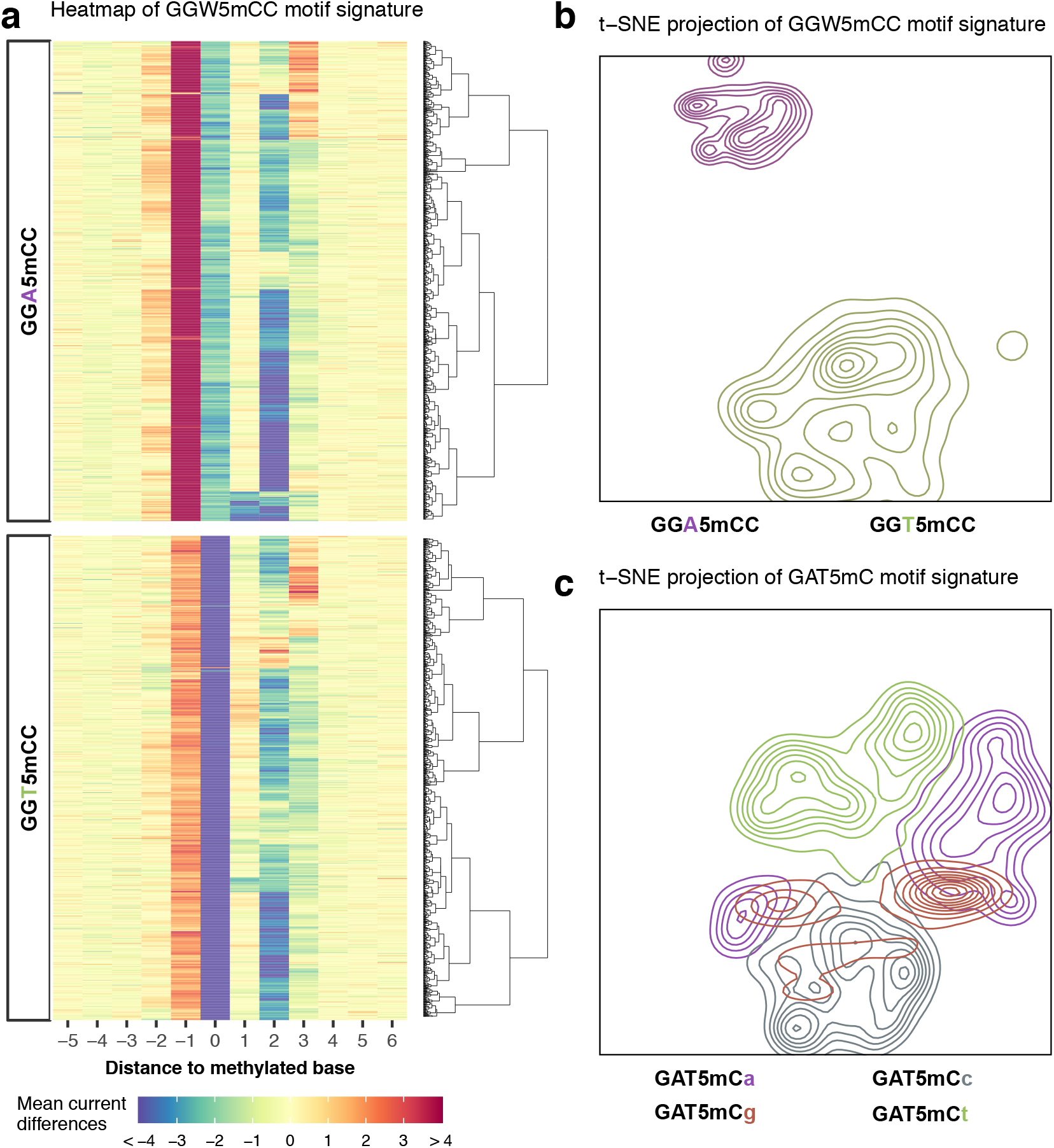
Local sequence context effect on motif signatures. (**a**) Sequence-dependent variation in current differences for GGW5mCC methylation motif occurrences. Current differences from violin plots of GGW5mCC in **Fig. 2a** were plotted as a heatmap with each row representing current differences flanking a methylation occurrence ([-5, +6] relative to methylation). GGW5mCC motif occurrences were split into two groups according to degenerated base (W=[A|T]) and ordered, within groups, using hierarchical clustering to highlight current difference patterns. (**b**) t-SNE projection of motif occurrences from **a** with cluster density displayed as relief. Clusters are colored according to degenerated base within the methylation motif. (**c**) Another example of sequence-dependent variation for GAT5mC motif occurrences with cluster density displayed as relief. Clusters are colored according to the first base following GAT5mC motif.

We explore other potential sources of variation by comparing general signal characteristics using t-SNE projection with different base caller versions (Albacore 1.1.0 vs 2.3.4), and different signal processing workflows (Tombo, and outlier removal). The base caller versions tested did not significantly change the signal characteristics (**Supplementary Fig. 3a**) and give consistent methylation detection performances (**Supplementary Fig. 3b**). Regarding signal processing workflow comparison, similar methylation signal properties were obtained when bacterial datasets were processed with Tombo compared to our pipeline, suggesting that no significant bias was introduced: motif signatures are similar (**Fig. 2a and Supplementary Fig. 4a**), methylation sites cluster by type (**Fig. 2c** and **Supplementary Fig. 4c**), by motif (**Fig. 2b and Supplementary Fig. 4b**), and also by local sequence context (**Fig. 3b and Supplementary Fig. 4d**). The outlier removal procedure reduces noise in the motif signatures (**Supplementary Fig. 3c-e**), and does not introduce significant bias in our signal processing as indicated by the high similarity with signal processed with Tombo (**Fig. 2a** and **Supplementary Fig. 4a**). Finally, we also confirmed that current differences obtained using a *de novo* assembled genome (*E. coli* at 200x; Methods) were consistent with the one obtained from matching reference genome ruling out the possibility that observed clustering pattern could be explained by an inaccurate reference genome (**Supplementary Fig. 3f,g**).

In summary, these analyses show that current differences induced by DNA methylation of the same type have great variation and heterogeneity in nanopore sequencing. This observation has important implications on methods development for nanopore sequencing based detection of DNA methylation. Specifically, it suggests that a broadly applicable method for methylation discovery is best trained using a comprehensive dataset with methylation motif diversity rather than a dataset of one or few specific motifs. This motivated us to develop the novel method that we will describe in the next section.

### A new method that enables *de novo* methylation typing and fine mapping

To account for the great signature diversity of methylation induced current differences across sequence contexts, we developed a novel method for the following two challenging tasks yet unaddressed by existing methods: 1) methylation typing, where the goal is to identify the type of DNA methylation, and 2) fine mapping, where the goal is to identify the position of the methylated base.

#### Methylation motif enrichment

Before introducing the novel method, we need to first describe the procedure we used for methylation detection and motif enrichment analysis building on existing methods^9, 14, 25^. In brief, 1) current levels are compared between native and WGA datasets for each genomic position (Methods); 2) p-values are combined locally with a sliding window-based approach followed by peak detection (Methods); 3) flanking sequences around the center of peaks are used as input for MEME motif discovery analysis (Methods). Overall, 45 of the total 46 well-characterized methylation motifs from seven bacteria were successfully re-discovered (**Supplementary Table 2**). The only undetected motif, GT6mAC from *H. pylori*, has much fewer occurrences (*i.e.* only 198 in the entire genome) than other 4-mer motifs (7169 occurrences on average). The motif discovery analysis also found six additional motifs not among the 46 well-characterized motifs (Methods, Supplement text), which were not included in subsequent analysis of our new method that is focused on methylation typing and fine mapping.

#### A novel method for de novo methylation typing and fine mapping

Although 45 of the 46 known motifs have already been re-discovered *de novo* in the above analysis, two critical additional features have yet to be defined: methylation type and methylated base within each motif. Despite the fact that the t-SNE analysis reveals a lack of a common signature for each methylation type and a large variation in current differences across different motifs of the same methylation type, it shows that DNA methylation events of the same type generally cluster well (**Fig. 2c**). We hypothesized that a classification model trained using diverse methylation types and motifs may serve as a reliable approach for categorizing *de novo* detected methylation into a specific methylation type.

In standard applications of classification models, both training and test samples need to be defined with respect to a consistent feature vector (*e.g.* current differences near methylated bases in our case). However, while both methylation type and methylation position are known for the well-characterized training samples (*i.e.* feature vectors can be consistently defined relatively to the methylated base for classifier training), features vector for the test samples cannot be aligned consistently because the methylated position is yet to be predicted. Essentially, methylation type classification and methylation fine mapping are coupled problems that need to be approached simultaneously.

Encouragingly, although the methylated base is not always at the center of the current differences, we did observe a relatively narrow window of no more than +/- 3 bp offsets from peak centers across the 46 well-characterized motifs (**Supplementary Fig. 5a**). This motivated us to design a novel multi-label classifier training strategy in which each well-characterized methylation occurrence is represented by multiple feature vectors with offsets relative to the known methylation position (+/- 3bp). Each methylation occurrence from a wide range of sequence context is learned 7 times by the classifier, each time using current differences at a specific offset from the methylated base. For a given test sample with unknown methylation type and unknown methylated position, the classifier will first use the center of current differences as an approximation of the methylated position and then predict the methylation type and the exact methylated position (Methods; **Fig. 4a-c**). This is the core design of our method that enables completely *de novo* methylation typing and fine mapping, which is critical for practical applications to unknown bacterial genomes.

**Figure 4:**
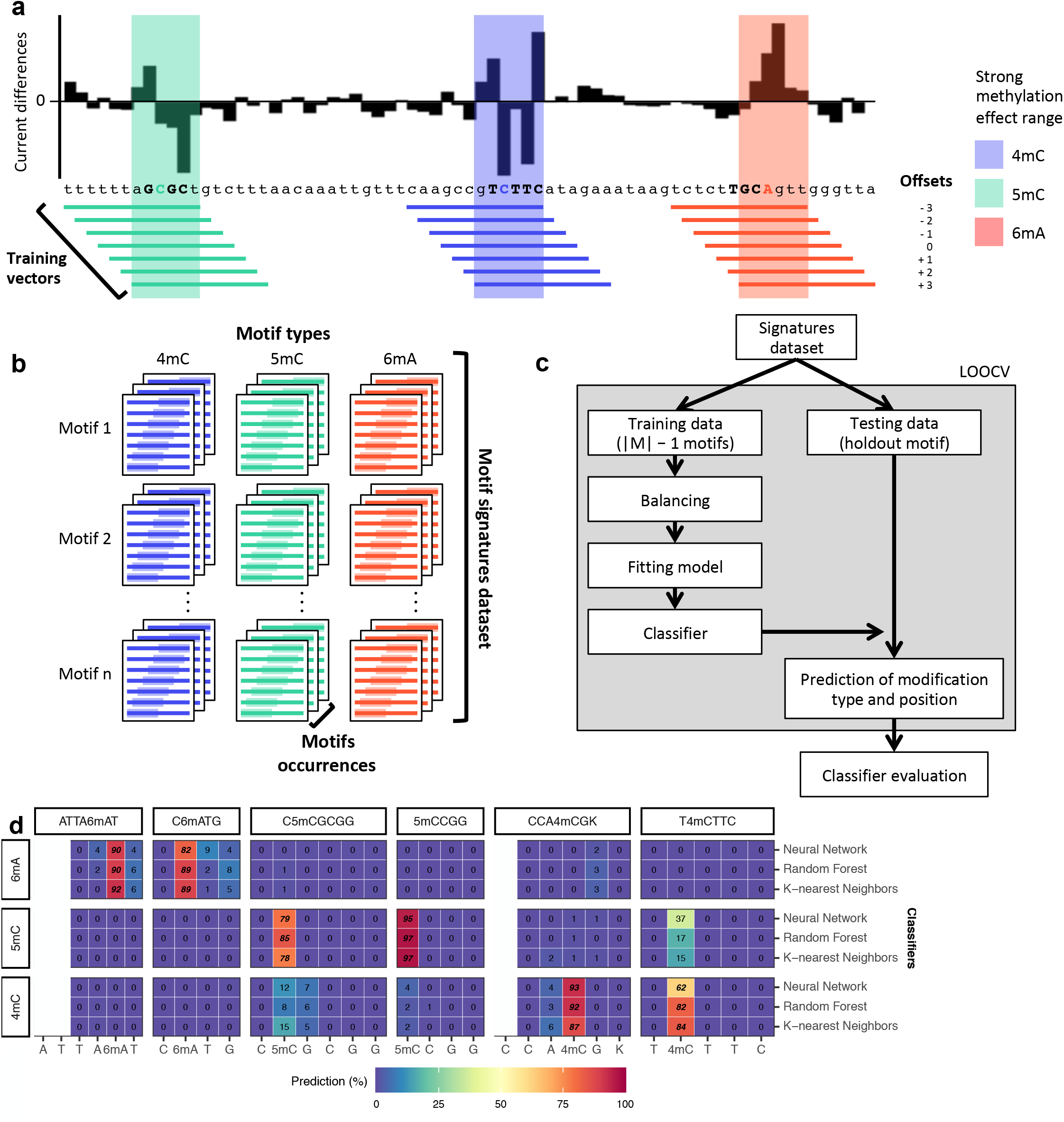
Classification and fine mapping of three types of DNA methylation. (**a**) Schematic representation of dataset building for classifier training. For each motif occurrence, we produced 7 training vectors of length 12 with +/- offsets from 0 to 3 position(s) relative to current differences core defined as [-2, +3] (**Supplementary Fig. 1a-c**). (**b**) Each training vector is labeled with the corresponding methylation type and offset used. They are then gathered into a large training dataset of current differences flanking 183,818 methylated bases from 46 distinct motifs (Methods). This dataset of current differences near the methylated base can be used to train classifiers. (**c**) Classifiers performances were evaluated using leave-one-out cross validation (LOOCV). (**d**) Subset of classifier evaluation results. Nine models were trained for each holdout combination to evaluate their performance for classifying holdout motifs. We classify every individual occurrence of each holdout motif and compute percentage of occurrences for each of the 21 labels using each classifier separately. Results for six selected motifs are shown as an illustration. Only within motif predictions are displayed, and the raw classification results for all motifs are presented in **Supplementary Fig. 6,7**. Filling colors correspond to percentage of occurrences classified to a specific class ranging from blue (0%) to red (100%). Prediction percentages of expected classes are displayed in italics and fine mapped methylated positions in each motif are displayed in bold.

A set of nine different classifiers was separately trained using current differences flanking known methylated bases following the offset strategy described above (Methods; **Fig. 4a-c; Supplementary Table 4**; **Supplementary Fig. 5b,d**). For classifier evaluation, we used leave-one-out cross validation (LOOCV) strategy where one motif is held out for testing while all the other 45 motifs are used for training. With all held out individual methylation sites belonging to a single methylation motif classified, predicted methylated type and position within motif was determined by using the consensus across tested occurrences (Methods). Overall results are largely consistent across the nine classifiers both in terms of accuracy for classifying individual methylation sites (**Supplementary Fig. 5c**) and methylation motifs, although k-nearest neighbors, random forest, and neural network had relatively better performances with at least 95.7% of motifs correctly typed and fine mapped (consensus accuracy of 97.8%; **Supplementary Fig. 6 and 7**).

Motif typing and fine mapping performances were further assessed on two independent bacterial samples: *N. otitidiscaviarum* and *T. phaeum*. All the 12 known methylation motifs were *de novo* re-discovered as well as accurately typed and fine mapped (**Supplementary Table 5**). In addition, we observed similar method performances when we used a *de novo* assembled genome (*E. coli* sample at 200x, 99.94% consensus accuracy) for motif detection (**Supplementary Fig. 3g**), and for the motif typing and fine mapping procedures (**Supplementary Fig. 3h**). We also evaluated the impact of genomic coverage using subsampled datasets of *H. pylori*, and observed improvement in motif enrichment (from 5x to 75x and then plateaued; **Supplementary Fig. 8a**; Supplementary text), as well as in motif typing and fine mapping across studied motifs (from 10x to 30x; **Supplementary Fig. 8c**).

In summary, we developed a new classification-based method that not only captures the complex variation of current differences across methylation types and motifs, but is also trained using a multi-label classification design that allow fine mapping of the methylated base. While we expect the method is highly reliable for *de novo* methylation typing and fine mapping for a methylation motif (97.8% consensus accuracy), we would like to note that the accuracy for individual methylation event varies dramatically across different motifs, ranging from 24% for G6mAGG, the only motif that was not accurately classified, to 99.5% for G5mCCGGC (median >76% for k-nearest neighbors, random forest, and neural network; **Supplementary Fig. 5c, 6, and 7**), which is consistent with the observation that motifs of the same methylation type can have different signatures (**Fig. 2c** and **Supplementary Fig. 2,3,8b**).

### Methylation discovery from microbiome and methylation-enhanced metagenomic analyses

Uncultured bacteria represent a significant proportion of the overall diversity of the bacterial kingdom and thus a great source for DNA methylation discovery. Therefore, we attempted to perform *de novo* methylation discovery and characterization from a mouse gut microbiome using nanopore sequencing. For microbes with fairly high abundance, metagenomic assembly often generates reasonably long contigs, which can be technically treated as individual genomes for methylation analysis using the procedure described in the last section. However, for microbes with relatively lower abundance, metagenomic assembly often results in fragmented genomes where contigs are short hence including only a limited number of occurrences of each motif, which makes methylation motifs discovery statistically underpowered if each metagenomic contig is examined separately.

Fragmentation related issues can be mitigated by using diverse binning methods intended to group related contigs together (species or strains level). Those methods encompass sequence composition features binning^26–29^, contig coverage binning^30–33^, as well as chromosome interaction maps^34–36^. Recent work by *Beaulaurier et al*. demonstrates that microbial DNA methylation can be exploited to enhance the grouping of metagenome contigs (*i.e.* methylation binning) using SMRT sequencing^37^. Instead of trying to discover precise methylation motifs from individual contigs, the methylation binning method presented in this recent work computes 6mA profiles (methylation scores for putative 6mA motifs) for each contig and then groups contigs together into bins based on methylation profiles similarities. We hypothesized that methylation binning of metagenomic contigs could be done using nanopore sequencing, which holds great promise due to its sensitivity for detecting all three types of common DNA methylations (4mC, 5mC, and 6mA) beyond the scope of work by *Beaulaurier et al*. that focused on 6mA alone^37^, especially because SMRT sequencing does not effectively detect 5mC.

We first developed a new methylation binning approach specifically for nanopore sequencing data considering the fundamental differences from SMRT sequencing (Methods; **Supplementary Fig. 9**). In a nutshell, several important technical steps needed to be developed for nanopore sequencing data because the current differences associated with each of the three types of methylation are spanning multiple events near methylated bases (**Fig. 2a**, **Fig. 3a**, and **Supplementary Fig. 1**) rather than confined to a single base for 6mA or 4mC as in SMRT sequencing. After prototyping and evaluation on a mock community (Supplement text; **Supplementary Fig. 10**), we applied the methylation method to new nanopore sequencing data of the same mouse fecal sample used in the SMRT sequencing-based study (MGM1; **Supplementary Table 6**). To summarize, after the *de novo* metagenome assembly using nanopore sequencing data only, we computed methylation feature vectors for a large set of candidate methylation motifs (n=210,176; Methods). Motifs with informational feature (*i.e.* significant current differences) were first selected based on large contigs, and methylation feature vectors were then computed in remaining contigs. Methylation feature vectors are then arranged in a methylation profile matrix, which is subjected to clustering analysis based on similarity among contigs (Methods). This initial automated binning resulted in ten bins (**Supplementary Fig. 11a**), which were further refined by per-bin motif detection followed by *de novo* motif guided binning (Methods; **Supplementary Fig. 11b-d**). The final methylation binning round of the mouse gut microbiome sample with nanopore sequencing data was performed using the 80 *de novo* detected methylation motifs (**Supplementary Table 7**) and revealed thirteen bins containing from 3 to 43 contigs in each (**Fig. 5a; Supplementary Fig. 11d**; **Supplementary Table 6**). The unique methylation profiles for each bin are displayed in **Figure 5c**. The method was further tested with a second microbiome sample, MGM2 (**Supplementary Table 6**), in which eleven bins with unique methylation profiles were identified (**Fig. 5b; Supplementary Fig. 12; Supplementary Table 9**).

**Figure 5:**
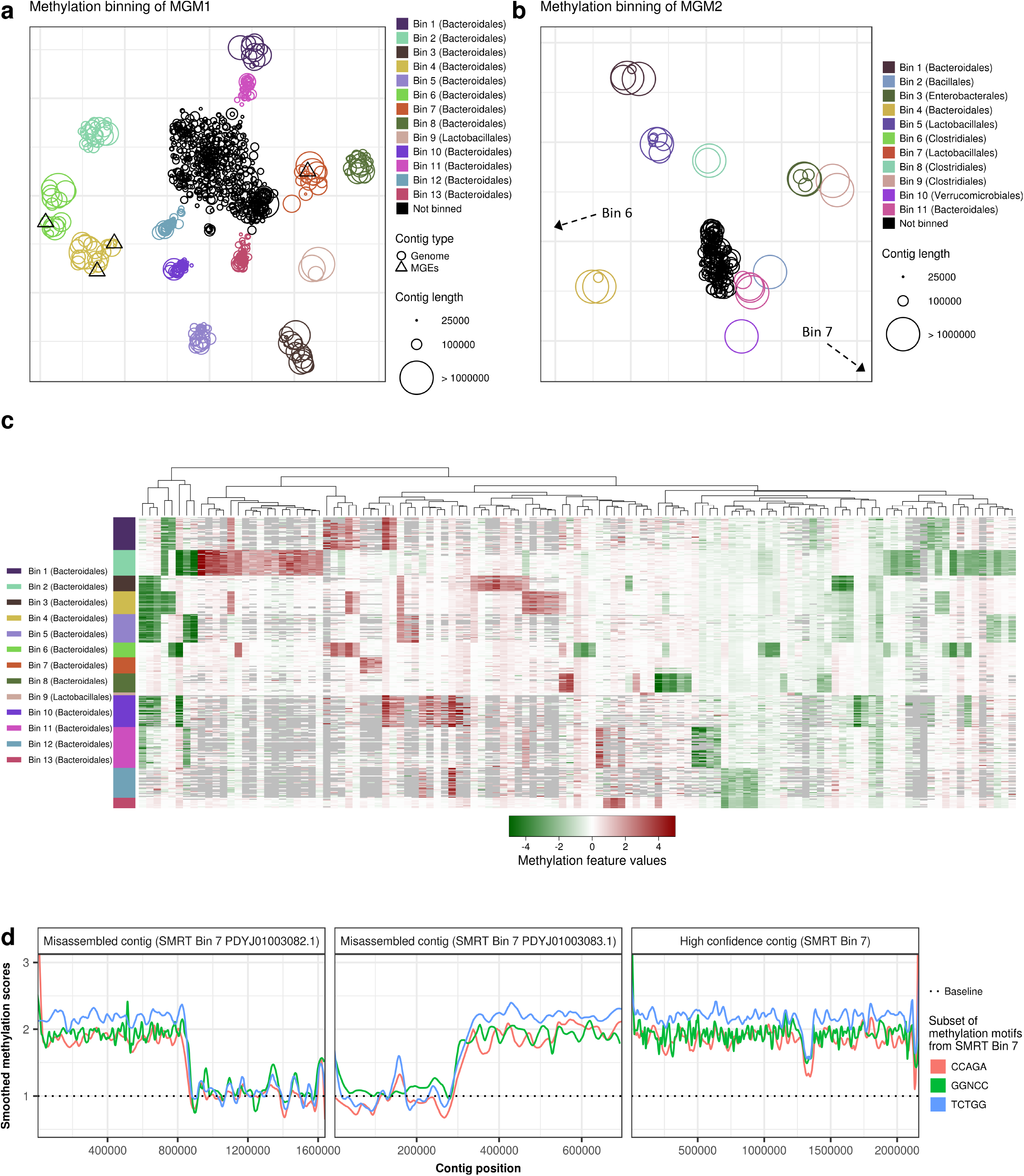
Methylation analysis of mouse gut microbiome samples. (**a**) Methylation binning of MGM1 metagenome contigs using *de novo* discovered motifs (after three rounds of binning followed by motif discovery; Methods, **Supplementary Fig. 9, Supplementary Fig. 11**). Methylation features computed from *de novo* discovered motifs are projected on two dimensions using t-SNE. Contigs are colored based on bin identities with point sizes matching contig length according to the legend. (b) Methylation binning of MGM2 metagenome contigs using *de novo* discovered motifs (after one round of binning followed by motif discovery; Methods, **Supplementary Fig. 9, Supplementary Fig. 12**). Methylation features computed from *de novo* discovered motifs are projected on two dimensions using t-SNE. Contigs are colored based on bin identities with point sizes matching contig length according to the legend. Non-zoomed plot (with visible Bin 6 and Bin 7) is presented in **Supplementary Fig. 12b**. (c) Heatmap representation of methylation feature values across binned contig from MGM1 sample. Methylation features were computed from all the *de novo* discovered motifs in MGM1 bins. Only the significant features with absolute values above 1.5 pA in the bin of origin (where the corresponding motif were discovered) were selected. Missing methylation features from contigs (less than 5 motif occurrences) are colored in grey. (d) Detection of misassemblies using methylation motif information along contigs. Left and middle panels: misassembled contigs mislabeled as Bin 7 in SMRT analysis (PDYJ01003082.1 and PDYJ01003083.1, contigs marked with an asterisk in **Supplementary Fig. 13a**. Right panel: an example of a properly assembled contig from Bin 7 (PDYJ01000763.1). We selected some *de novo* detected motifs from Bin 7, and scored their methylation sites along the three contigs. Methylation scores were then smoothed using locally estimated scatterplot smoothing and displayed with one color per motif. Smoothed methylation scores are consistent in the contig from the right panel, but not in the misassembled contigs shown in the two left most panels. A switch of methylome occurs near 800 kbp and 300 kbp in the left two panels respectively, supporting the existence of misassemblies (detailed in **Supplementary Fig. 14a,b**).

We also performed the analysis using the SMRT metagenomic assemblies reported in the recent study^1^ to ease the comparison between nanopore sequencing and SMRT sequencing (Methods). Through a bin-level comparison, bins from nanopore sequencing data closely matched those from SMRT sequencing data, and none of the nanopore sequencing bins contained misclassified contigs (**Supplementary Fig. 13a** and **Supplementary Table 10**). Consistent between the two technologies, methylation binning effectively separated the multiple Bacteroidales species that are usually hard to distinguish from each other due to their highly similar genome sequence composition and abundance^37^.

Through the methylation binning analysis, 80 methylation motifs (62 unique recognition sequences) were discovered from the thirteen bins from MGM1 sample (**Supplementary Table 7**). We applied the methylation typing and fine mapping method trained in the previous section to these 80 methylation motifs and compiled classification results from the consensus across k-nearest neighbors, random forest, and neural network classifiers. Methylation typing and fine mapping predictions are consistent with the motif recognition sequences for 64 motifs: 18 are identified as 6mA, 45 as 5mC and 1 as 4mC (**Supplementary Table 11**). The rarity of 4mC motifs is consistent with a reanalysis of SMRT sequencing data from the recent study^37^, which also confirmed every 6mA motif discovered with our method. The *de novo* detection of a large number of 5mC motifs is very encouraging because previous large-scale bacterial methylome studies were almost exclusively based on SMRT sequencing, which is known to be ineffective for detecting 5mC methylation. However, not every 6mA motif found with SMRT sequencing was detected in the analysis of nanopore sequencing data. The missing ones are mostly bipartite 6mA motifs, which are usually not frequent and thus more challenging to detect using nanopore sequencing. This is probably due to the diffuse nature of current differences around 6mA (**Fig. 2a** and **Supplementary Fig. 1**) in contrast to the highly specific signal right on top of 6mA in SMRT sequencing.

We further attempted to link mobile genetic elements (MGEs) to their host genome based on their methylation profiles. Using the SMRT metagenomic assembly with *de novo* discovered methylation motifs, we were able to bin 11 of the 19 annotated MGEs from this microbiome sample according to their methylation profiles (five plasmids and six conjugative transposons; **Supplementary Fig. 13b; Supplementary Table 12**), while nine were binned with the SMRT analysis^37^. With eight MGEs binned as with SMRT analysis and three newly binned MGEs, nanopore sequencing increased MGEs linking potential compared to SMRT methylation binning likely owing to its better sensitivity to 5mC motifs. From our nanopore-only *de novo* metagenome assembly, fewer MGEs were identified (eight), although similar results were obtained in terms of linking MGEs to their host genomes, *i.e.* four out of the eight MGEs identified were binned correctly (**Fig. 5a**).

In addition to contig binning, we hypothesized that the microbial DNA methylation pattern can also be used to discover misassembled contigs. In a nutshell, the methylation pattern is expected to be largely consistent across different regions of an authentic metagenomic contig. Following this rationale, we discovered two contigs from SMRT sequencing based metagenomic assembly of the MGM1 sample (marked by an asterisk in **Supplementary Fig. 13a**) that both show inconsistent intra-contig methylation status (**Fig. 5d**). By comparing methylation patterns from methylation motif sets from the other bins, we found that the contigs in question are chimeric contigs both representing Bacteroidales species (**Supplementary Fig. 14**, Bin 2 and Bin 7). This is consistent with the previous examination of coverage uniformity and contamination through single-copy gene count^37^, confirming that those contigs annotated as Bin 7 were misassembled by HGAP2 combining parts of Bin 2 and Bin 7 genomes. Generally, this analysis highlights the benefit of incorporating DNA methylation status (ideally all three types: 6mA, 4mC and 5mC), which not only help better distinguishing microbes species but also help assess contig homogeneity revealing eventual misassemblies, an application particularly useful for the characterization of complex microbiome samples.

## Discussion

In this work, we developed a novel method for *de novo* discovery (methylation typing and fine mapping) of three forms of bacterial DNA methylation, namely 4mC, 5mC, and 6mA. We demonstrated that it enables the much-needed *de novo* characterization of unknown bacterial methylomes from both individual bacteria and microbiome samples. Our comprehensive motif profiling and analysis showed that different methylation motifs of the same methylation type could differently impact current levels captured in nanopore sequencing. This observation has important implications for nanopore sequencing based detection of DNA methylation confirming that a rich collection of methylation sequence context is necessary to develop broadly applicable computational methods for methylation discovery, which we achieved through aggregation of a diverse assortment of methylation motifs from bacteria. As increasing number of researchers start to employ nanopore technology for microbial genome sequencing, we expect our new method to be widely used for methylation discovery from bacteria.

As we attempted to use the novel method to directly detect DNA methylation and discover methylation motifs from a microbiome, we demonstrated two valuable utilities of DNA methylation analysis by nanopore sequencing for helping to characterize metagenomes. First, we developed a novel approach for methylation binning of metagenomic contigs and linking of MGEs to host genomes building on the method reported for SMRT sequencing data^37^ and designing multiple technical procedures addressing the unique properties of nanopore sequencing, *i.e.* the diffuse nature of current differences around methylation events in contrast to the highly specific signal right on top of 6mA and 4mC events in SMRT sequencing. Second, we demonstrated that examining the methylation pattern along assembled metagenomic contigs could help identify chimeric contigs due to metagenomic misassemblies. The long reads from nanopore sequencing holds great promise in metagenome analysis, so we expect our new method to further help researchers exploit microbial DNA methylation for high-resolution metagenomic and MGE analyses.

While both SMRT sequencing and nanopore sequencing have great promise of direct detection of DNA methylation without the need for chemical conversions, there has not been an in-depth comparison between the two methods in terms of their advantages and disadvantages. In this aspect, our comparative analysis over the metagenomic contigs binned by methylation motifs detected by the two technologies from the same microbiome sample provided important insights. First, while 5mC is very challenging to detect using SMRT sequencing, nanopore sequencing provides reliable 5mC detection. The large number of 5mC motifs discovered from the mouse gut microbiome sample using nanopore sequencing suggests the prevalence and diversity of 5mC motifs could have been largely underestimated in the >2,000 bacterial methylome analyses that were almost exclusively based on SMRT sequencing^1^. Second, we found that multiple long and rare methylation motifs well detected by SMRT sequencing in the metagenome analysis were missed by nanopore sequencing, which can be explained by the current differences associated with each of the three types of methylation diffusion to multiple flanking bases in contrast to the fairly high IPD ratios confined to a single methylation site (4mC or 6mA) for SMRT sequencing^2, 20–23^. Collectively, these comparisons suggest that SMRT sequencing and nanopore sequencing have their own strengths and limitations, hence the two technologies are expected to complement each other in various applications.

In this work, we focused on bacterial methylomes of individual microbes and microbiomes, and we expect the method to be highly reliable for *de novo* methylation typing and fine mapping for methylation motifs. For individual methylation sites, we would like to highlight that the accuracy of the current method for methylation typing and fine mapping varies across different motifs, which calls for development of more accurate methods in future work. Also, because motif occurrences are almost 100% methylated in bacterial methylomes, the current method did not attempt to estimate the fraction of molecules methylated at each genomic position. In practice, existing tools such as Tombo allow the estimation of partial methylation at individual methylation sites, thus are complementary to our method.

Last but not least, although the current study was focused on three types of DNA methylation, similar design could be extended for the detection of additional forms of DNA methylation (5hmC, 5fC and 5caC) as well as other forms of DNA chemical modification such as the various forms of DNA damage^38, 39^, and possibly diverse forms of RNA modifications^40, 41^ owing to the unique promise of nanopore technology for direct RNA sequencing^42–44^.

## Acknowledgements

We thank Yimeng Kong and Mi Ni for providing helpful feedback for early versions of this manuscript. The work was supported by a seed fund from Icahn Institute for Genomics and Multiscale Biology (G.F.), and by R01 GM128955 (G.F.) from the National Institutes of Health. G.F. is a Hirschl Research Scholar by Irma T. Hirschl/Monique Weill-Caulier Trust, and a Nash Family Research Scholar. This work was also supported in part through the computational resources and staff expertise provided by the Department of Scientific Computing at the Icahn School of Medicine at Mount Sinai.

## Competing interests

A.T. and G.F. are inventors of two US Provisional patent applications (62/860,952 and 62/851,205) that describe the methods in this manuscript.

## Author contributions

G.F. conceived and supervised the project. A.T. and G.F. designed the methods. A.T. developed the software package for all the proposed computational analyses. A.T., E.A.M., and X-S.Z. conducted the experiments. A.T. and G.F. analyzed the data, and wrote the manuscript with inputs and comments from all co-authors.

## Methods

### Software and data availability

Software of the novel methods and a detailed tutorial with supporting data are available at http://github.com/fanglab/nanodisco. All sequencing data generated in this study will be made available upon publication.

### Samples collection and DNA extraction

A set of nine bacteria was rationally selected using a previous study^5^ and REBASE^19^ to provide a large diversity of methylation motifs, in particular for the less frequent 4mC and 5mC methylation motifs: *Bacillus amyloliquefaciens* H, *Bacillus fusiformis* 1226, *Clostridium perfringens* ATCC 13124, *Escherichia coli* K-12 substr. MG1655 ATCC 47076, *Methanospirillum hungatei* JF-1, *Helicobacter pylori* JP26, *Neisseria gonorrhoeae* FA 1090, *Nocardia otitidiscaviarum* NEB252, and *Thermacetogenium phaeum* DSM 12270.

*B. amyloliquefaciens* H, *B. fusiformis* 1226, and *N. otitidiscaviarum* NEB252 DNA samples were obtained from New England Biolabs (NEB, Ipswich, MA). Those for *C. perfringens* ATCC 13124, *M. hungatei* JF-1, *H. pylori* JP26, *N. gonorrhoeae* FA 1090 and *T. phaeum* DSM 12270 were obtained from the Human Health Therapeutics Research Area at National Research Council Canada, the Department of Microbiology, Immunology, and Molecular Genetics at University of California Los Angeles, the Department of Medicine at New York University Langone Medical Center (NYUMC), the University of Oklahoma Health Sciences Center, and the Department of Biology at the University of Konstanz (Germany), respectively. Finally, we obtained *E. coli* K-12 substr. MG1655 ATCC 47076 directly from the American Type Culture Collection (ATCC, Manassas, VA).

The adult mouse gut microbiome DNA samples (MGM1 and MGM2) were obtained from the Department of Medicine at NYUMC. MGM1 DNA sample was extracted from the fecal pellets used in the SMRT sequencing study^37^ while MGM2 DNA sample comes from fecal pellets of the same mouse after antibiotic treatment with tylosin. Fecal DNA extraction was performed using QIAamp DNA Microbiome Kit (QIAGEN, Hilden, Germany) followed by cleanup with DNA Clean & Concentrator – 5 elution buffer (ZYMO Research, Irvine, CA) and final elution in 10 mM Tris-HCl, pH 8.5, 0.1 mM EDTA.

### Library preparation and sequencing

The quality of input DNA was controlled with Nanodrop 2000 and concentration measured using Qubit 3.0 (Thermo Fisher Scientific, Waltham, MA). Native libraries were prepared following 1D Genomic DNA by ligation protocol (SQK-LSK108; version GDE_9002_v108_revT_18Oct2016) with minor modifications described below. Whole genome amplification samples were prepared using REPLI-g Mini Kits (QIAGEN, Hilden, Germany) according to the protocol with 12.5 ng of input DNA and 16 h incubation. Next, WGA samples were treated with T7 endonuclease I (NEB) to maximize nanopore sequencing yield according to ONT documentation. WGA libraries were prepared following Premium whole genome amplification protocol from T7 step (version WAL_9030_v108_revJ_26Jan2017) with minor modifications described below. Bacteria (other than *E. coli* and *H. pylori*) and mouse gut microbiome DNA samples, native and WGA, were RNase A treated (FEREN0531, Thermo Fisher Scientific) then fragmented at 8 kbp with g-TUBEs (Covaris, Woburn, MA) to homogenized DNA fragments lengths increasing accuracy of input DNA molarity calculation to maximize yields. Final fragment length distributions were determined using Bioanalyzer 2100 (Agilent Technologies, Santa Clara, CA). Samples were sequenced on R9.4 and R9.4.1 flow cells (**Supplementary Table 3** and **6**).

*E. coli* and *H. pylori* libraries (native and WGA) were prepared without fragmentation or Formalin-Fixed, Paraffin-Embedded (FFPE) DNA repair. *E. coli* and *H. pylori WGA* input DNA was increased to 3 μg in T7 step with 20 min incubation. Remaining steps were performed according to corresponding ONT protocol and final libraries sequenced on 3 flow cells with a maximum of two consecutive runs per flow cell. Flow cells were washed between runs using the Flow Cell Wash Kit (EXP-WSH002) from ONT. An additional WGA was produced for *H. pylori*, and referred to as independent WGA. Sequencing of native and WGA libraries for *E. coli* and *H. pylori* generated from 289 to 2630x genomic coverage but were down sampled at 200x to more accurately represent common yield targets.

DNA samples for the additional bacteria (*B. amyloliquefacien*, *B. fusiformis*, *C. perfringens*, *M. hungatei*, *N. gonorrhoeae*, *N. otitidiscaviarum, and T. phaeum*) were pooled in equimolar quantity for library preparation. Pooling possibility was confirmed by mapping mock nanopore reads datasets generated using Nanosim^45^ (version 1.0.0; simulator.py linear -r <path_to_fasta> -c <error_model> -o <path_output> -n 50000 -- min_len 200 --max_len 50000 using the *E. coli* error model provided by the authors on 03/23/17) on the combined references and verifying accurate separation of reads into genome of origin. Any reads mapping on more than one genome were discarded from all the analysis presented in our study, independently of the mapping type. Native and WGA library preparations were performed using aforementioned ONT protocol and sequenced on separate flow cells (**Supplementary Table 3**). Sequencing of native and WGA generated datasets with coverage ranging from 65 to 297x.

Finally, mouse gut microbiome libraries (MGM1 and MGM2) were generated according to the One-pot ligation protocol for Oxford Nanopores libraries (dx.doi.org/10.17504/protocols.io.k9acz2e) including the FFPE DNA repair step with exception for the room temperature incubation times that were increased from 10 to 20 minutes. 300 fmol of input DNA were used in FFPE DNA repair steps. Native and WGA libraries were sequenced on separate flow cells for 48 h (**Supplementary Table 6**).

### Nanopore sequencing signal processing

Nanopore sequencing reads are base called using ONT Albacore Sequencing Pipeline Software (version 2.3.4). Reads are mapped to corresponding references using BWA-MEM (version 0.7.15 with *–x ont2d* option)^46^. The following steps are performed using R (version 3.5.3)^47^. Reads are separated by strand according to the initial alignment (package Rsamtools; version 1.34.1)^48^, and both groups are processed as forward strand reads by mapping reverse strand reads on the reverse complement of the reference genome using BWA-MEM. Supplementary and reverse strand alignments are then filtered out with samtools (version 1.3; flags 2048 and 16)^49^. Next, events are associated to genomic positions according to alignment coordinates from reads and expected current levels with Nanopolish *eventalign* (version 0.11.0)^9^. Event levels are normalized across reads by correcting signal scaling and shifting. Both normalization factors are computed for each read by fitting events level to ONT 6-mer model (nanopolish configuration file r9.4_450bps.nucleotide.6mer.template.model) using robust regression (rlm function). Event level outliers are removed using Tukey’s fences methods based on interquartile range (IQR=1.5) for each genomic position. Finally, mean event current differences (pA) were computed by comparing event levels between native sample (maintained methylation state) and WGA sample (essentially methylation free) at each genomic position for both strands separately. This metric is simply referred to as current differences in our manuscript. Associated p-values from two-sided Mann-Whitney U test are also computed (*wilcox.test* function) which was proposed in Stoiber et al.^14^. Only genomic positions with sufficient coverage are considered in later analysis (min_cov=5).

### Motif enrichment analysis

DNA methylation affects nanopore sequencing signal at multiple positions around the methylated base (**Fig. 2a** and **Supplementary Fig. 1a-c**)^3^ meaning detection of methylated sites can be reinforced by combining information from consecutive genomic positions. As in Stoiber et al., consecutive p-values are combined with Fisher’s method (*sumlog* function) in sliding windows (5 bp) smoothing statistical signal along the genome^14^. It combines the methylation related signal near methylated bases and reduces signal noises from spurious genomic positions. Resulting smoothed statistical signals form peaks near methylated positions. Detected peaks are ranked according to their smoothed p-value and the top 2000 peaks are then selected for motif discovery. An alternative strategy is to randomly sample peaks from more than the top-2000 positions (Supplementary text). Corresponding genomic sequences are then extracted (22 bp) and used as input for *de novo* motifs discovery with MEME software (version 4.11.4; parameters: -dna -mod zoops -nmotifs 5 -minw 4 -maxw 14 -maxsize 1000000)^25^. Selection of region of interest based on combined p-values followed by motif detection using MEME was initially proposed in a preprint by Stoiber et al.^14^. However, we enhanced the motif discovery potential by closely integrating MEME in our pipeline as described in next paragraphs.

Running time for motif discovery with MEME rapidly increases with size of the sequence dataset to such extent that we had to limit the number of input sequences used. To address this constraint, we adopt a repeated procedure of back and forth between peak detection and motif discovery steps. For each pass, a limited number of input sequences are analyzed with MEME and motifs achieving a sufficient confidence (E-value <= 10^-30^) are reported. After each motif discovery step, peaks explained by discovered motifs, whose corresponding genomic sequence contains at least one of the *de novo* detected motifs, are removed making it possible to discover less frequent motifs and ones with weaker signals. This repeated procedure is adapted for detecting any number of methylated motifs while decreasing processing time.

Raw motifs called by MEME were further refined by leveraging current difference information. For each motif reported by MEME, we generated a list of mutated motifs by introducing a substitution (one substitution at a time; analysis of GATC will give 12 mutated motifs: AATC, CATC, TATC, GCTC, GGTC, GTTC, GAAC, GACC, GAGC, GATA, GATG, GATT). We then compute each mutated motif signature (see Motifs typing and fine mapping) with associated scores representing total divergence from nonmethylated signature (sum of absolute average current differences).

### Parameter tuning for signal processing and motif detection

To assess our methods performance for *de novo* motif discovery and tune parameters, we evaluated the enrichment of MEME input sequences for expected motifs as the chosen smoothed p-value threshold varies. Method development and choice of default parameters was guided by evaluating various metrics including Precision-Recall (PR), Receiver Operating Characteristic (ROC) curves and area under curves (AUC). We used the following two comparisons to define contingency table classes: native versus WGA, and independent WGA versus WGA. True positives (TP) and false negatives (FN) are respectively defined as motif occurrences with or without signal peak above threshold in native versus WGA. False positives (FP) are genomic regions without motifs and with signal peak above threshold in native versus WGA as well as motif occurrences with signal peak above threshold in independent WGA versus WGA. Finally, true negatives (TN) are defined as genomic regions without motifs and without peak above threshold in native versus WGA as well as motif occurrences without peak above threshold in independent WGA versus WGA. State of motif occurrences were defined whether a peak was detected above the chosen threshold in a 22 bp window encompassing expected methylated base of motif occurrences. For genomic regions devoid of motif, those were split in 22 bp consecutive units, and used as FP and TN with similar status definition. Performances were computed on the first 500 kbp of the reference genome only. When comparing performances for *de novo* detection between individual motifs, we took into consideration variation in frequencies (*i.e.* a rare motif will be more difficult to detect). Therefore, in order to make the evaluation more generally applicable, we fixed the ratio of positive regions (22 bp windows from motif occurrences in native versus WGA) over all queried regions to one third by random subsampling, effectively avoiding variation in frequencies across the set of *H. pylori* motifs.

Using the aforementioned method, we evaluated parameter performances for *de novo* methylation detection for the following steps or parameters: read mapping, event current normalization, outlier removal (**Supplementary Fig. 3c,d**), statistical test, p-value combining function, smoothing window size, and peaks window size. We also evaluated the impact of coverage by subsampling at 10 depths ranging from 5x to 200x as well as the impact of motif frequency and the motif specific context (*i.e.* how methylation type and sequence context affect detection potential; **Supplementary Fig. 8**).

### Validation of methylation motifs used for classification

*E. coli* and *H. pylori* were sequenced with SMRT sequencing in order to confirm 4mC and 6mA methylation motifs using the RS_Modification_and_Motif_Analysis protocol from SMRT Analysis Server (v2.3.0). Methylation status summaries for the remaining bacterial species (modifications.csv and motif_summary.csv files) were obtained from the U.S. Department of Energy Joint Genome Institute and NEB. We confirmed effective methylation of 4mC and 6mA motifs individually by checking if IPD ratio consistently peaked on expected methylated bases. Finally, REBASE annotation was used as a gold standard for 5mC motifs. Methylation motifs with an ambiguous status (*e.g.* weak or partial IPD ratio peaks) or not reported in REBASE annotation were not used for the classifier training and the performance evaluation.

### Motif typing and fine mapping

For each bacterial genome, we list methylated genomic positions from each strand based on motif recognition sequences. Methylated positions in close proximity are discarded to avoid introducing unwanted complexity (at least 22 bp apart, each strand considered independently as current signal is strand specific). Ambiguous motifs are removed from downstream analysis (see Validation of methylation motifs used for classification in Methods). We extract current differences in [- 10 bp, + 11 bp] range relative to methylated base positions. Each occurrence is labeled with genome of origin, recognition sequence, methylation type, methylation position within motif, and genomic coordinates. This dataset constitute our methylation motif signatures for motif typing and fine mapping, while we use a range of [- 6 bp, + 7 bp] to examine the variation of current differences across different DNA methylation types and motifs. Note that for *de novo* detected methylation motif and refinement function, signatures are generated considering every position in the motif as potentially methylated, which produced a longer signature not necessarily centered on the methylated base.

The training dataset for classification is generated from methylation motif signatures to permit labeling of methylation type and position within motifs simultaneously (**Fig. 4a**). For each vector of current differences from a methylated site, we generate 7 smaller vectors, lengths 12, offseted by one position so that each of them still contains the [- 2 bp, + 3 bp] range relative to the methylated base. In other words, those 7 vectors contain current differences from the [- 2 bp, + 3 bp] range with up to 3 additional position(s) before or after (*i.e.* [- 5 bp, + 6 bp] +/- 0 to 3 bp). Each of those vectors is labeled with the type of DNA methylation from corresponding motifs as well as corresponding offset used (from - 3 to + 3) resulting in 21 different labels (7 offsets x 3 types DNA methylation).

For the testing datasets, methylated base position is unknown and current difference vectors cannot be defined in the same way. However, methylated base position can be approximate by computing the center of current differences from a motif signature. For that, we average absolute current differences from a motif signature using a sliding window of length 5 and the position with the largest variation is used as an approximation of methylation position within the motif (**Supplementary Fig. 5a**). In practice, approximations are not further than 3 bp from the methylated position meaning that the vectors of current differences centered on those approximations will match one type of vector offset used for training because they are generated with - 3 to + 3 bp offsets.

Prior to any model fitting, the training dataset is balanced by random sampling to contain a similar number of vectors for each label in order to avoid bias toward the more common methylation type. Classifier hyperparameters (**Supplementary Table 4**) were tuned on the balanced training dataset containing all motifs using repeated 10-fold cross validation (n=3) with balanced accuracy (mean and standard deviation) as the main metric. Robustness of chosen hyperparameters was confirmed by comparing performances from three classifiers (k-nearest neighbors, random forest, and neural network) when using parameters either tuned on a dataset containing all motifs (as described above) or a dataset only containing *H. pylori* motifs only. Both sets of hyperparameters gave similar results when tested on a dataset without *H. pylori* motifs (**Supplementary Fig. 5d**).

Classifier performance evaluation was performed using leave-one-out cross validation strategy (LOOCV) by holding out current differences vectors from one motif and training on remaining vectors (from all motifs except one). The resulting model is then used to predict the label of held out vectors from the tested motif. The LOOCV strategy simulates models behavior when faced with an unseen motif signature. For testing, we only used the set of vectors corresponding to the approximated methylation position found as described previously. Predicted methylated base type for a motif is defined using consensus across all tested motif occurrences. As for methylated base position, the classifier prognosticates the offset between the approximated methylation position chosen as input and the predicted methylation position, which is then converted into a position within tested motifs.

### Nanopore sequencing based *de novo* assembly

Genome assembly for *E. coli* was performed using Canu^50^ (version 1.8; *-nanopore-raw genomeSize=4.7m overlapper=mhap utgReAlign=true*) with the native nanopore reads (200x dataset). Next, we generated the genomic consensus with Racon^51^ (version 1.3.3; default parameters) to correct raw contigs, and correct contig ends using nucmer^52^ (version 4.0.0beta; *--maxmatch –nosimplify* and *show-coords -lrcTH*) to identify and trim remaining overlaps. Then, we polished the assembly consensus using Nanopolish^9^ (version 0.11.0; *variants --min-candidate-frequency 0.1* for five rounds) with the native nanopore reads. Finally, we performed another round of polishing with Nanopolish using nanopore WGA reads (methylation free) to correct remaining assembly error caused by DNA methylation signal in the native reads (same parameters for five rounds).

### Metagenome methylation binning

While methylation motif detection could be performed as for individual bacteria, metagenome assemblies often result in many contigs from multiple organisms with various lengths making individual contig analysis lacking power. Instead, we propose to first bin contigs with similar methylation profiles then perform the motif detection. Nanopore sequencing native and WGA datasets are processed in the same way as for individual bacteria (except that supplementary alignment were conserved) generating current differences alongside metagenome contigs using the nanopore sequencing-only *de novo* metagenome assembly.

*De novo* metagenome assemblies for MGM1 and MGM2 were performed using Flye^53^ (version 2.4.2; *--meta –nano-raw –genome-size 100M*) with the native nanopore reads. Next, the metagenome consensus was computed using Racon^51^ for four consecutive rounds (default parameters). Then, the resulting metagenome assemblies were polished using Nanopolish^9^ with first the native, then with the WGA nanopore reads (*variants --min-candidate-frequency 0.1* for five rounds with each set of reads).

For a candidate motif, an associated methylation feature vector is computed by averaging current differences from aggregated occurrences on a metagenomic contig (**Supplementary Fig. 9**). Unlike well-characterized methylation motifs, the methylated position in a candidate motif is unknown. Therefore, we consider every position in motifs as potentially methylated by including all potentially affected current differences in the methylation feature vector calculation. For a motif of length k, we compute a methylation feature vector of length k + (2 + 3), which corresponds to the length of current differences that are possibly affected by a methylated base in a k-mer motif (the core current differences is defined as [- 2 bp, + 3 bp] range flanking a methylated base, **Supplementary Fig. 1**). This procedure results in a methylation feature vector of average current differences of length k + 5 representing a motif methylation status for a contig. This step represents a major difference from SMRT sequencing based methylation binning method where a single methylation score is generated for a motif on 37 a contig^37^.

The next step is to create a methylation profile matrix comprising methylation feature vectors for each motif of interest in each metagenomic contig, which will be used for methylation binning (**Supplementary Fig. 9**). A set of 210,176 candidate motifs is generated according to common structures (4-, 5-, and 6-mers, as well as bipartite motifs with 3 to 4 bp specificity part separated by 5 to 6 bp gaps). In order to select motifs of interest, an initial round of motif evaluation is performed on a subset of longer contigs (100 kbp using nanopore sequencing *de novo* assembly) with sufficient coverage (10x; **Supplementary Fig. 10**) with the rationale that results will have a higher statistical power. Uninformative methylation features are filtered out by discarding the ones with small absolute current difference values across the initial contig set (< 1.5 pA; chosen based on our mock metagenome analysis) as well as the ones computed from fewer than 20 motif occurrences. Next, we additionally filtered out uninformative methylation features from bipartite motifs by removing methylation feature vectors with fewer than two significant features across the initial contig set (significant if current difference >= 1.5 pA) to account for the longer vector and generally lower motif frequency. Finally, methylation features from bipartite motifs that overlap with any remaining 4 to 6-mer motifs are also discarded. The resulting list of informative methylation features is then evaluated in each contig of the metagenome assembly to construct a methylation profile matrix (**Supplementary Fig. 9**). This two-step approach effectively reduces the initial research space on the set of large contigs speeding up the analysis, and reduces noise by only considering methylation features selected from contigs with higher statistical power. The resulting methylation profile matrix (significant methylation features computed across all contigs) is then processed using t-SNE dimensionality reduction method to visualize contig clusters (**Supplementary Fig. 9**). Missing methylation features and ones computed from fewer than 5 motifs occurrences are set to small random pseudovalues in the [- 0.2, + 0.2] range (reducing correlation from missing methylation features; random number generation seeds are set at 2, 3, and 4 for MGM1, MGM2, and the SMRT assemblies respectively). Small contigs are not considered for methylation binning (<25 kbp for the nanopore sequencing *de novo* assembly analysis), and remaining ones are weighted according to their length. Weighting factors are defined as quotient of contig length divided by 50,000 and capped at a percentage of the number of remaining contigs to avoid extreme imbalance (only contigs with coverage >= 10x for both native and WGA are weighted). We set the capping value at 5% for metagenome with high diversity (large number of metagenome contigs, MGM1) and 10% for simpler metagenome (<500 contigs, MGM2). Finally, bins are defined after t-SNE dimension reduction using DBSCAN (package dbscan version 1.1-4), an automated clustering method, with additional manual annotation of visible bins that can be missed by DBSCAN.

The analysis using the SMRT metagenome assembly (GCA_002754755.1) is performed as described previously using thresholds of 500 kbp and 10x of coverage for initial methylation feature selection (contigs from Bin 3, Bin 4, and Bin 9 are not covered sufficiently due to the use of a different DNA extraction kit than the SMRT study). Contigs smaller than 10 kbp are not considered.

Motif detection from bins is performed the same way as for individual bacteria. With *de novo* detected motifs, methylation feature vectors used for binning are not filtered, keeping the full-length methylation feature vectors. Missing methylation features from individual contigs are handled as described previously and contigs are also weighted. We performed three rounds of binning and motif detection for MGM1 (**Supplementary Fig. 11**), while one was sufficient for MGM2 (**Supplementary Fig. 12**). Confirmation of *de novo* discovered motifs in MGM1 sample (potential 6mA and 4mC motifs) from nanopore sequencing analysis were realized with per bin motif detection from SMRT sequencing data using the SMRT portal pipeline (RS_Modification_and_Motif_Analysis.1).

Binning focused on associating mobile genetic elements (MGEs) to host genome (**Supplementary Fig. 13b**) was performed using metagenome reference from the SMRT study where binned contigs were replaced by per-bin reassemblies^37^. MGEs contigs from the nanopore-only *de novo* metagenome assemblies were identified according to the alignment of MGEs sequences from the SMRT study using minimap2 (version 2.15; *- ax asm20*)^54^.

### Detection of metagenome contigs misassemblies

The rationale is to examine the consistency of methylation signal for a motif across different occurrence of the motif along a metagenomic contig. For every single motif occurrence, we calculate a score by taking the average of absolute current differences from six consecutives positions with the most perturbation. Then, these individual scores are averaged using a sliding window across the contig to examine the continuity. Motif occurrences from both strands are used in this analysis. However, if a motif occurrence overlaps with another motif site being examined (<15 bp) then both are discarded.

## Notes

https://github.com/fanglab/nanodisco

