## Supplemental Text for "Discovering and exploiting multiple types of DNA methylation from individual bacteria and microbiome using nanopore sequencing"

### Supplementary figures captions

**Supplementary Figure 1:** General statistics of motif signatures. (**a**) Distribution of current differences are shown for all confident motifs altogether as well as average absolute differences and associated standard deviations near methylated bases ([- 10, + 11]). (**b**) Same as **a** with distinction between DNA methylation types. (**c**) Same as **a** but for individual methylation motifs.

**Supplementary Figure 2:** Systematic examination of three main DNA methylation types with nanopore sequencing. (**a**) t-SNE projection of isolated methylation motif occurrences separated per motif. The same dataset as **Fig. 2b** was used with occurrences colored per motif. Other motifs are colored in grey. (**b**) Same as **a**, but grouped by methylation type.

**Supplementary Figure 3:** Nanopore sequencing signal processing variable. (**a**) Comparison of current differences across methylation occurrences between datasets base called with Albacore 1.1.0 and Albacore 2.3.4 illustrated by projection with t-SNE from for 46 well-characterized motifs (**Supplementary Table 2**). Each dot represents one isolated motif occurrence colored by base caller versions. For each motif occurrence, current differences from 22 positions near methylated bases ([- 10 bp, + 11 bp]) were used. (**b**) Performance for *de novo* methylated site detection between datasets base called with Albacore 1.1.0 and Albacore 2.3.4. We evaluated individual motif occurrences detection using Precision-Recall curves for *H. pylori* at 75x coverage. Precision-Recall curves and area under the curves (AUC) were computed as described in the Method section. Only confident *H. pylori* motifs were considered for the evaluation. (**c**) Comparison of current differences across methylation occurrences (same as **a**) between datasets produced with or without outlier removal step (Methods). (**d**) Performance for *de novo* methylated site detection (similar than **b**) with datasets produced with or without outlier removal step. (**e**) Variation of current differences across methylation occurrences without outlier removal step as illustrated by motif signatures from three motifs (AG4mCT, GGW5mCC, and GCYYG6mAT). For each motif, current differences near methylated bases ([- 6 bp, + 7 bp]) from all isolated occurrences are plotted with conservation of relative distances to methylated bases. Distributions of current differences for each relative distance are displayed as a violin plot. Current differences axis is limited to -8 to 8 pA range. (**f**) Comparison of current differences across methylation occurrences (same as **a**) with *E. coli* datasets (200x) produced using either the reference genome or the *de novo* assembly (Methods). (**g**) Performance for *de novo* methylated site detection in *E. coli* datasets (200x) using either the reference genome or the *de novo* assembly. (**h**) Performance of methylation motif typing and fine mapping on *E. coli* datasets (200x) produced using either the reference genome or the *de novo* assembly. Only results for k-nearest neighbors, neural network, and random forest are displayed.

**Supplementary Figure 4:** Systematic examination of three main types of DNA methylation with nanopore sequencing and Tombo signal processing. (**a**) Variation of current differences computed with Tombo across methylation occurrences as illustrated by motif signatures from three motifs (AG4mCT, GGW5mCC, and GCYYG6mAT). For each motif, current differences near methylated bases ([- 6 bp, + 7 bp]) from all isolated occurrences are plotted with conservation of relative distances to methylated bases. Distributions of current differences for each relative distance are displayed as a violin plot. Current differences axis is limited to -8 to 8 pA range. (**b**) Variation of current differences computed with Tombo across methylation occurrences as illustrated by projection with t-SNE from for 46 well-characterized motifs (**Supplementary Table 2**). Each dot represents one isolated motif occurrence colored by methylation motif. For each motif occurrence, current differences from 22 positions near methylated bases ([-10 bp, + 11 bp]) were used. (**c**) Similar to **b** but colored by DNA methylation type with additional processing to reveal cluster density indicated by relief. (**d**) Local sequence context effect on motif signatures. Sequence-dependent variation in current differences computed with Tombo for GGW5mCC methylation motif occurrences. t-SNE projection of motif occurrences from GGW5mCC in **a** with cluster density displayed as relief. Clusters are colored according to degenerated base within the methylation motif.

**Supplementary Figure 5:** Additional information for classification of methylation motif occurrences. (**a**) Approximation of DNA methylation position in three motifs (AGCT, GCYYGAT, and GGWCC). Signal strength is computed using a sliding window alongside motif signature to choose the best vector positioning to use for classification. (**b**) Flowchart description of procedure for classifier training and novel motifs dataset annotation. (**c**) Boxplot of overall prediction accuracy in LOOCV evaluation for each classifier. Classifiers are ordered by average accuracy. (**d**) Effect of hyperparameters on classification accuracy. Boxplot of overall prediction accuracy in LOOCV evaluation with classifiers trained on all motifs except the ones from *H. pylori*. Hyperparameters were either tuned on *H. pylori* motifs only ("Alt. HP") or on all motifs ("Main HP").

**Supplementary Figure 6:** Classification and fine mapping of three types of DNA methylation (part 1). Similar to **Fig. 4b** with full set of prediction results for a subset of methylation motifs. Filling colors correspond to percentage of occurrences classified to a specific class ranging from blue (0%) to red (100%). Greyed out prediction correspond to out of motif position. Blank columns correspond to within-motif positions without prediction. Prediction percentages of expected classes are displayed in italic and selected predictions based on consensus are displayed in bold.

**Supplementary Figure 7:** Classification and fine mapping of three types of DNA methylation (part 2). See **Supplementary Fig. 6**.

**Supplementary Figure 8:** Evaluation of motif enrichment with Precision-Recall curves. (**a**) Effect of coverage on *de novo* methylated site detection. We evaluated individual motif occurrences detection using Precision-Recall curves (PR curves) for *H. pylori*. Studied datasets with coverage ranging from 5x to 200x were generated by random subsampling of native and WGA datasets. Precision-Recall curves were generated as described in the Method section. We considered only confident *H. pylori* motifs for evaluation. (**b**) Precision-Recall curves summarizing the detection performance at 75x coverage of individual methylation sites for each motif in *H. pylori* with adjusted frequency (Methods). (**c**) Performance of methylation motif typing and fine mapping on datasets with genomic coverage subsampled at 10x, 15x, 20x, and 30x. Only results for k-nearest neighbors, neural network, and random forest are displayed.

**Supplementary Figure 9:** Schematic representation of methylation feature vectors computation and methylation binning of contigs. The computation of methylation features and the building of the methylation profile matrix is described in the method.

**Supplementary Figure 10:** Theoretical performance of methylation binning on mock microbiomes. Impact of contig coverage and contig length was assessed by generating mock metagenomes from individual bacteria datasets subsampled at 6 depths (genomic coverage of 5x, 10x, 15x, 20x, 30x, and 50x) with contig lengths varying from 5 kbp to 50 kbp (Supplementary text). Cluster silhouette coefficients were computed for each species (n=7) at each contig length and depth combination.

**Supplementary Figure 11:** Detailed methylation analysis of MGM1 sample. (**a**) Methylation binning using automated methylation features selection (without precise methylation motif discovery; Methods). Methylation features are projected on two dimensions using t-SNE. Contigs are colored per bin defined using DBSCAN, with point sizes matching contig length according to the legend. Two bins with the same methylation motifs were manually merged into Bin 4. (**b**) Methylation binning using *de novo* discovered motifs on each bin found in **a** (Methods). Methylation features computed from *de novo* discovered motifs are projected on two dimensions using t-SNE. Contigs are colored per bin defined using DBSCAN except Bin 11, which was manually defined. (**c**) Methylation binning using *de novo* discovered motifs on each bin found in **b**. Contigs are colored per bin defined using DBSCAN except for Bin 13, which was manually defined. (**d**) Methylation binning of MGM1 metagenome contigs using *de novo* discovered motifs (after three rounds of motif discovery (same as **Fig. 5a**).

**Supplementary Figure 12:** Detailed methylation analysis of MGM2 sample. (**a**) Methylation binning using automated methylation features selection (without precise methylation motif discovery; Methods). Methylation features are projected on two dimensions using t-SNE. Contigs are colored per defined bin with point sizes matching contig length according to the legend. Bin 1, 3, 4, and 5 were defined using DBSCAN. The other bins are composed of one or two contigs and were manually defined after *de novo* methylation motif discovery. (**b**) Methylation binning using *de novo* discovered motifs on each bin found in **a** (Methods). Methylation features computed from *de novo* discovered motifs are projected on two dimensions using t-SNE. Contigs are colored per bin as described in **a**.

**Supplementary Figure 13:** Methylation analysis of MGM1 sample with SMRT metagenome assembly. (**a**) Automated methylation binning of MGM1 metagenome contigs (without precise methylation motif discovery). Methylation status of common motifs (n=210,176) was screened across large contigs (>=500 kb) through computation of methylation feature vector (Methods, **Supplementary Fig. 9**). Informative motifs were selected and their status evaluated across remaining contigs. Resulting methylation features are projected on two dimensions using t-SNE. Contigs are colored based on bin identities assigned in the SMRT study[37] with point sizes matching contig length according to legend. Our binning identified two contigs originally identified as Bin 7 clustered separately from the main bin (contigs marked with an asterisk) suggesting that they have a different methylation profile than remaining Bin 7 contigs, which was not observed with SMRT analysis. Contigs marked with an asterisk are used as example for misassembly detection in **Fig. 5d**. (**b**) Methylation based association of MGEs to host genomes. Annotation of potential MGEs was obtained from the per-bin reassemblies from the SMRT study[37]. Genomic contigs are colored by bin of origin with point sizes matching their length. Some contigs are now binned within different clusters than their bin of origin likely because the original contigs were chimeric. Note that many contigs from Bin 7 reassemblies are affected because of the two major misassembled contigs identified.

**Supplementary Figure 14:** Detection of misassemblies in Bin 7 contigs from methylation motif signal. Identification of contamination origin for the two contigs mislabeled as Bin 7 (PDYJ01003082.1 and PDYJ01003083.1, marked with an asterisk in **Supplementary Fig. 13a**). We scored occurrences from methylation motifs found in each bin separately and smoothed signal along misassembled contigs (Methods). Scores from motif occurrences overlapping Bin 7 motifs were removed. Scores from Bin 2 motifs are consistently high in the second half of contig PDYJ01003082.1 and first half of contig PDYJ01003083.1 suggesting contamination originated from Bin 2 genomic sequences.

### Supplement Text

#### Nature of nanopore sequencing signal from Oxford Nanopore Technologies

Raw nanopore signal corresponds to electric current level (pA) sampled at 4000 hz across the nanopore while a DNA strand is transferred from one compartment to the other in a 450 bp.s-1 ratcheting motion. Higher order of signal structure, called events, consists in consecutive signal level corresponding to multiple measures of current for a specific relative position of the DNA strand inside the pore. The initial signal processing performed by the base caller, Albacore (version 1.1.0), detects those consecutive events and translates them into a nucleotide sequence.

#### *De novo* methylation motif detection with MEME

Running time for motif discovery with MEME increases with the number of input sequences therefore we limited the number of input sequences used to 2000 with the current implementation and parameters used. Furthermore, we observed that, with some genomes, top peaks (based on smoothed p-value) could be enriched in specific motifs combination (*i.e.* motifs in close proximity) preventing MEME from discovering individual motifs in favor of the specific motifs combination. This is due to larger than average smoothed p-value happening when two motif occurrences are near each other, which affect current in a broader genomic region. This phenomenon was observed for genomes with multiple frequent motifs. To limit this bias when observed, we provide an option to randomly select sequences among peaks above a threshold resulting in more than 2000 peaks, effectively avoiding the enrichment of specific motif combination.

#### Additional information for methylation motif validation

Our *de novo* methylation motif detection analysis also discovered six motifs not included in the confident list. Two motifs were detected in *H. pylori* (*i.e.* GGWTAA and GGWCNA, likely 6mA on sixth position) but the analysis of SMRT sequencing data suggest that they are partially methylated. Two additional motifs were found in *N. gonorrhoeae.* One of them is GTANNNNNCCC, likely modified by the MTase of GT6mANNNNNCTC, but SMRT data shows weak methylation signal suggesting that the motif is partially methylated (not methylated in all reads). The other one is TCACC, a 5mC methylation motif according to our classification (*i.e.* T5mCACC), which would explains why it was not detected with SMRT sequencing analysis. Finally, YGGCCR and WGGCCW were discovered in *B. fusiformis* and *C. perfringens* respectively. While both were expected to be the non-degenerated methylation motifs GG4mCC, SMRT sequencing data analysis also suggests that the others subsets of motifs (non-YGGCCR or non-WGGCCW) were not fully methylated explaining our results.

Other unconfident methylation motifs were found only with SMRT sequencing. In *H. pylori*, we listed three unconfident motifs (*i.e.* CTGG6mAG, CCTCT6mAG, and STA6mATTC) with weak signals suggesting that they were false discovery or partially methylated motifs (not methylated in all reads), thus not suitable for our study. However, we also found a methylation motif in *N. gonorrhoeae* with strong SMRT sequencing signal (*i.e.* CC6mACC) while little to no sign of methylation are visible with ONT analysis (*i.e.* no perturbation in average current differences near motif). It's unclear if this particular methylation motif is not detected because ONT method is not sensitive to change in nucleotide (between A and 6mA) in CCACC sequence context or because it's not methylated in our *N. gonorrhoeae* sample thus it was not used in our analysis.

Note that all the ambiguous motifs mentioned in this section were treated as potential methylation motifs when removing overlapping signal in order to avoid possible compound effects. However, they were ignored from all analysis.

#### Limiting factor for methylation motif detection

Genomic coverage strongly affects methylation motif detection ability with substantial improvement in motifs enrichment up to 75x in *H. pylori* with 20% to 90% of motif detected by increasing coverage from 5x to 75x (**Supplementary Fig. 8a**). Overall, 75x (37.5x per strand) is sufficient to detect 100% and 90% of motifs in *E. coli* and *H. pylori* respectively. In addition, we observed variation in enrichment across motifs even when variation in motifs frequency was accounted for (**Supplementary Fig. 8b**). Motif specific performances depend on the amount of current perturbation introduced by the methylation compared to the non-methylated signal. For example, the G6mAGG motif signature displayed weak current differences and was not detected for *H. pylori* dataset at lower coverage (<20x). At lower coverage, undetected motifs can display a clear signature although not sufficient to be enriched enough to detect them. Finally, in practice, bacterial methylation motifs have various frequencies in genomes sometimes independent of their complexity, which seems to be a limiting factor for their detection (e.g. GT6mAC in *H. pylori*). Note that while methylation motif signatures represent how DNA methylation affect ionic current in a specific genomic context during sequencing, some of their characteristics depend on the data processing method used (*e.g.* base caller, reads mapper, event aligner, and normalization). We expect that methylation motif detection performance will increase with improvement of nanopore sequencing preprocessing methods, notably for base calling and signal alignment to a reference sequence.

#### Approximation of methylated position from motif signature

Our current method for approximating methylated position within *de novo* detected motifs relies on the identification of the center of the motif signature. However, other educated guesses could be made based on motif signature and refining plots, which would permit reducing the DNA methylation position research space. First, main current differences are in the [- 2 bp, + 3 bp] range from the methylated base meaning that for bipartite motifs one could ignore part of the motif depending on which specificity subunit is aligned with current differences. Similarly, this could be done for long motifs if current differences are at one of the motif extremities. This phenomenon is indirectly used in our approximation approach. Second, motif signatures display important variation when the methylated base is close to non-fixed bases, *i.e.* next to a degenerated base or near motif extremities. This strategy was not used in the current implementation.

#### Mock microbiome from individual bacteria

In order to define motif selection procedure for contig methylation binning, we constructed a mock metagenome assembly from our individual bacteria reference genomes (n=7). Reference genomes were fragmented following mouse gut metagenome contig length distribution from previous SMRT study[1]. Nanopore sequencing native and WGA datasets subsampled at a coverage of 50x were then mapped on the mock metagenome assembly and processed similarly to individual genomes to generate current differences and associated U test p-values (Methods). Possible methylation motifs from the initial set (n=210,176) are scored for long contigs (>=500 kbp) according to the procedure described in Methods. Rules for methylation motif features selection were defined to enrich the final list in known methylation motifs from bacteria in the mock community. Only genomic positions with 10x coverage were used in both scoring steps.

We applied the following cutoff on methylation features: minimum absolute current differences (1.5 pA), minimum number of motif feature occurrences per confident contigs (20), minimum number of significant features in bipartite motifs (2), and discard overlapping motifs (bipartite motif explained by 4 to 6-mers motifs). Any motif features satisfying those requirements are scored in remaining contigs. Missing methylation features and those computed from fewer than 5 motifs occurrences were set to small random pseudovalues in the [-0.2, 0.2] range to reduce correlation from missing methylation features.

We further evaluated the impact of coverage on methylation binning performance by generating a mock metagenome from individual datasets subsampled at 6 depths (genomic coverage of 5x, 10x, 15x, 20x, 30x, and 50x). Similarly, we also evaluated the impact of contig length on binning by creating a mock metagenome with contigs of length varying from 5 kbp to 50 kbp. For each combination of the contig lengths and sequencing depths, we computed methylation features for the expected methylation motifs with genomic frequency above one per 10 kbp and performed t-SNE based dimensionality reduction followed by quantitative evaluation using average cluster silhouette coefficients[2]. Cluster silhouette coefficients were computed for each species (n=7) at each contig length and depth combination. We observed a clear improvement of average cluster silhouette coefficients at 10x compared to 5x for all contig length considered (**Supplementary Fig. 10**). Furthermore, as expected, as contig length increases, the performance of methylation binning also increases because more motif instances make the methylation feature estimation more accurate, reducing noise and making common methylation profiles easier to group together.

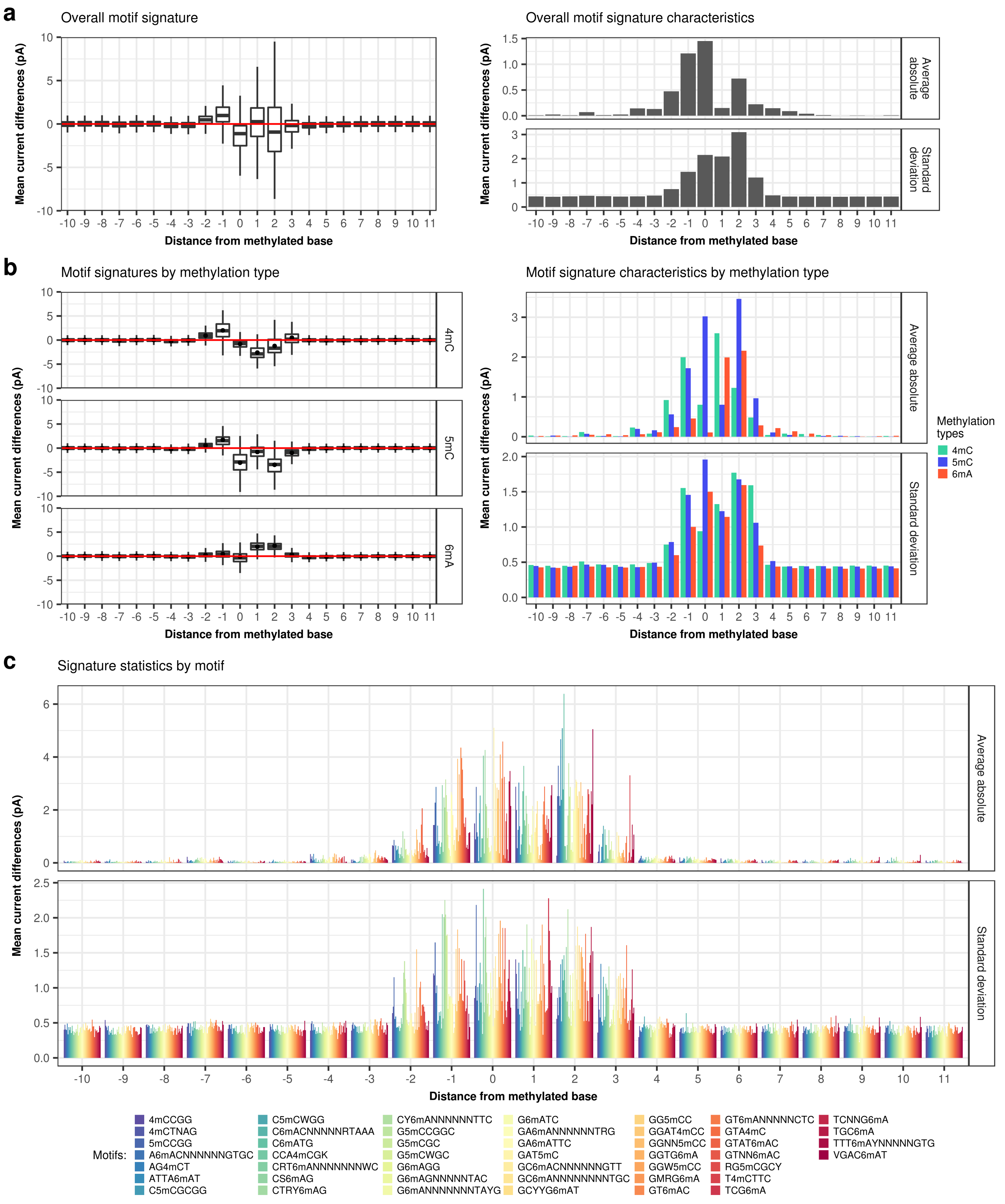


Supplementary Figure 1

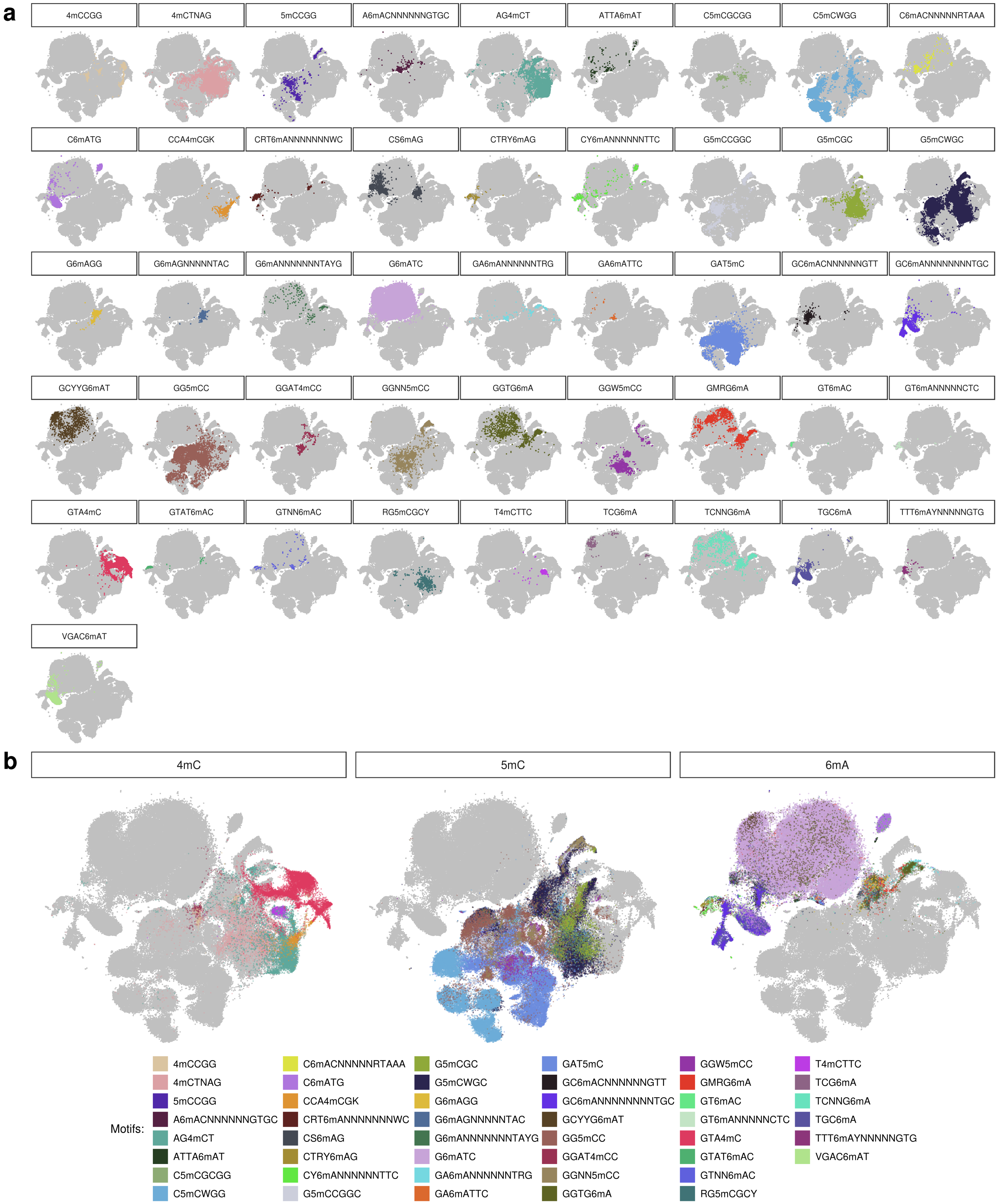


Supplementary Figure 2

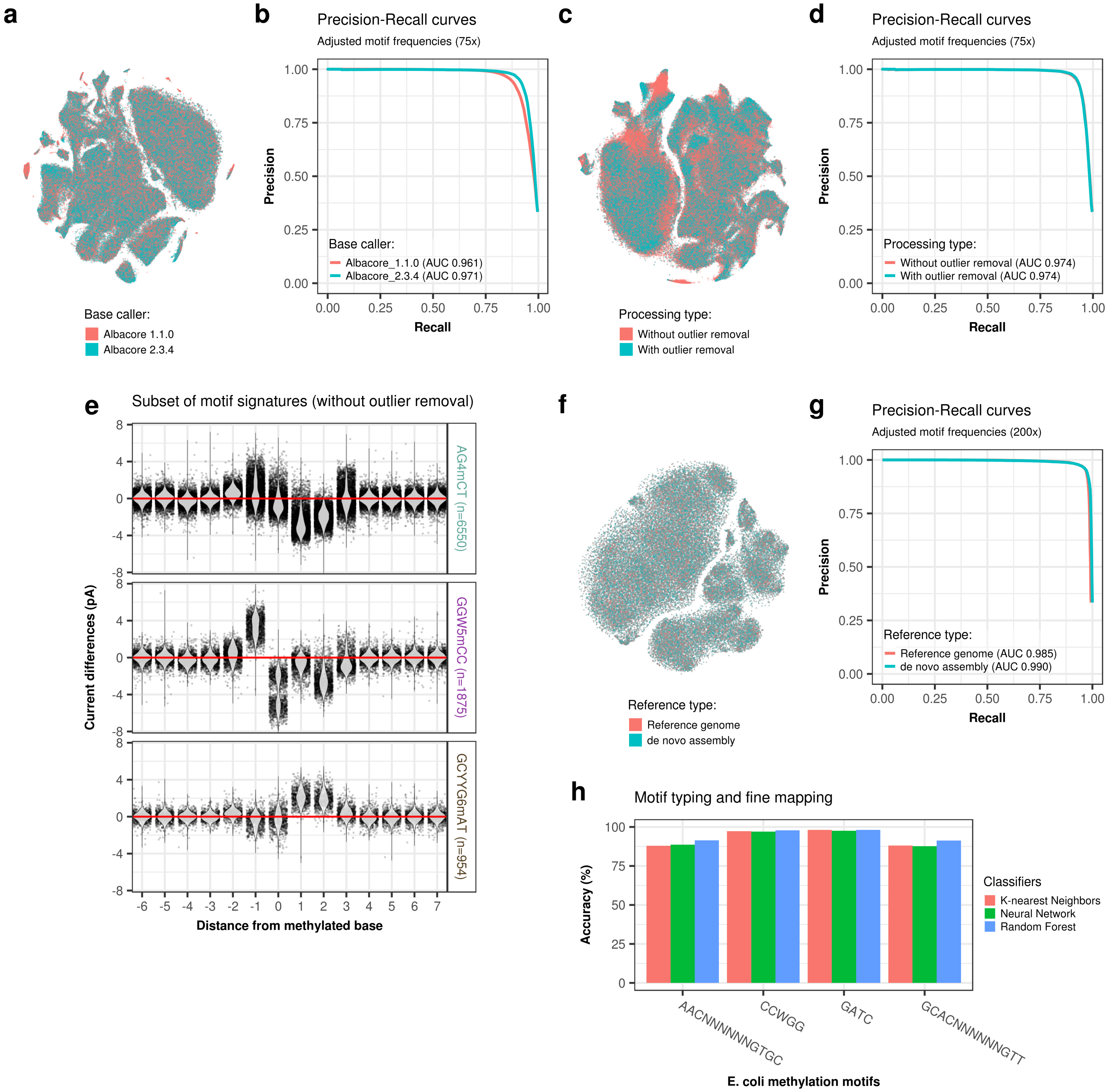


Supplementary Figure 3

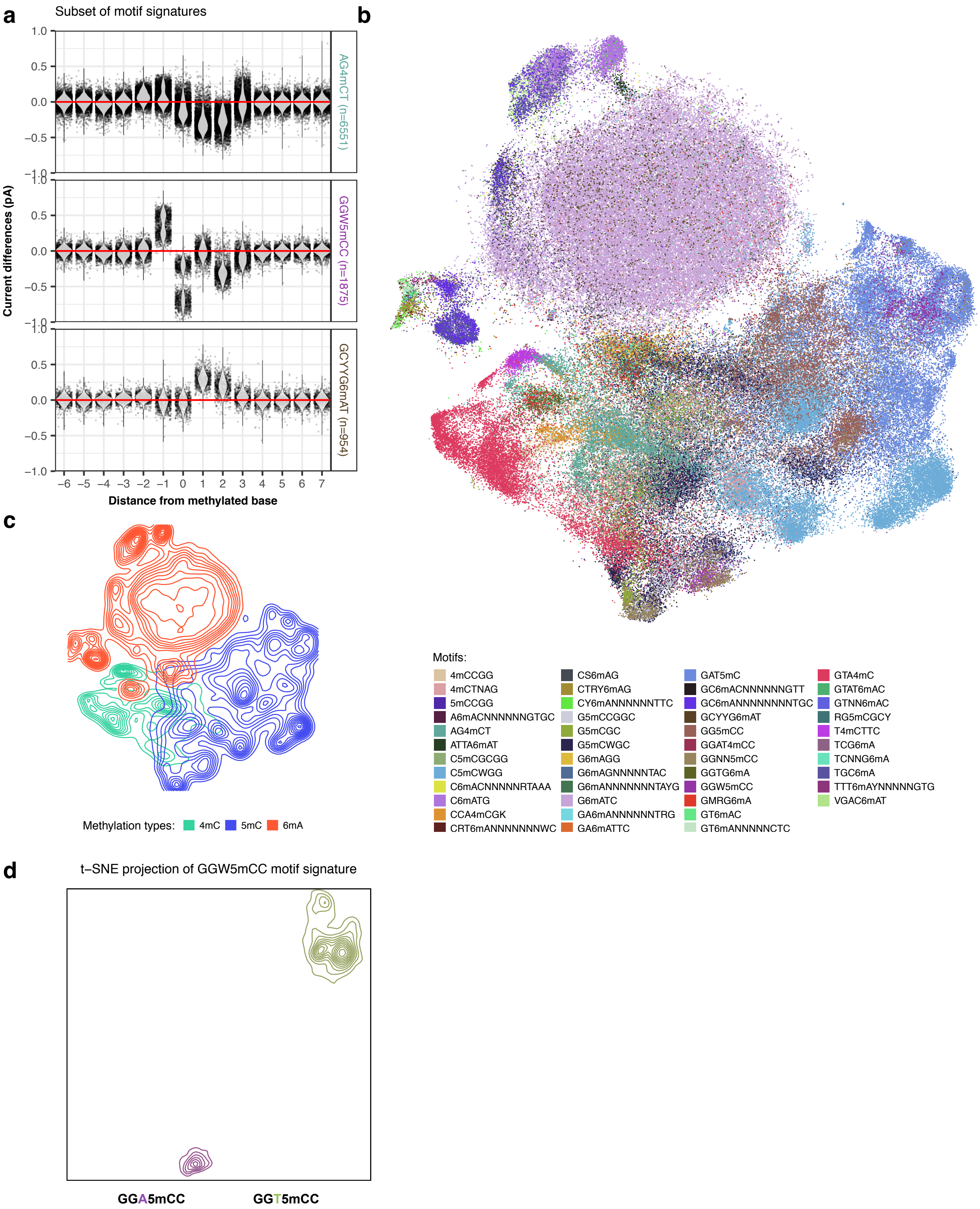


Supplementary Figure 4

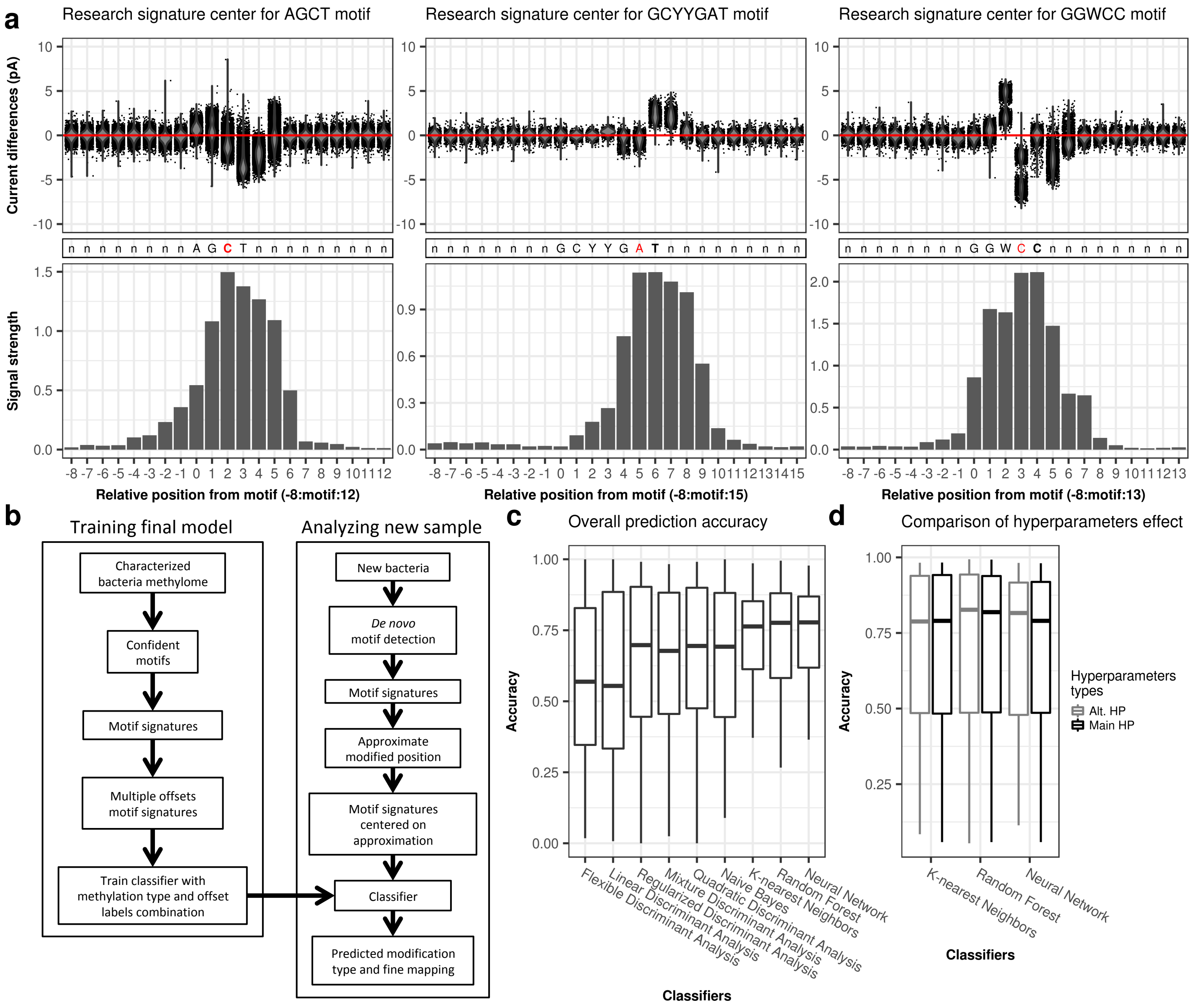


Supplementary Figure 5

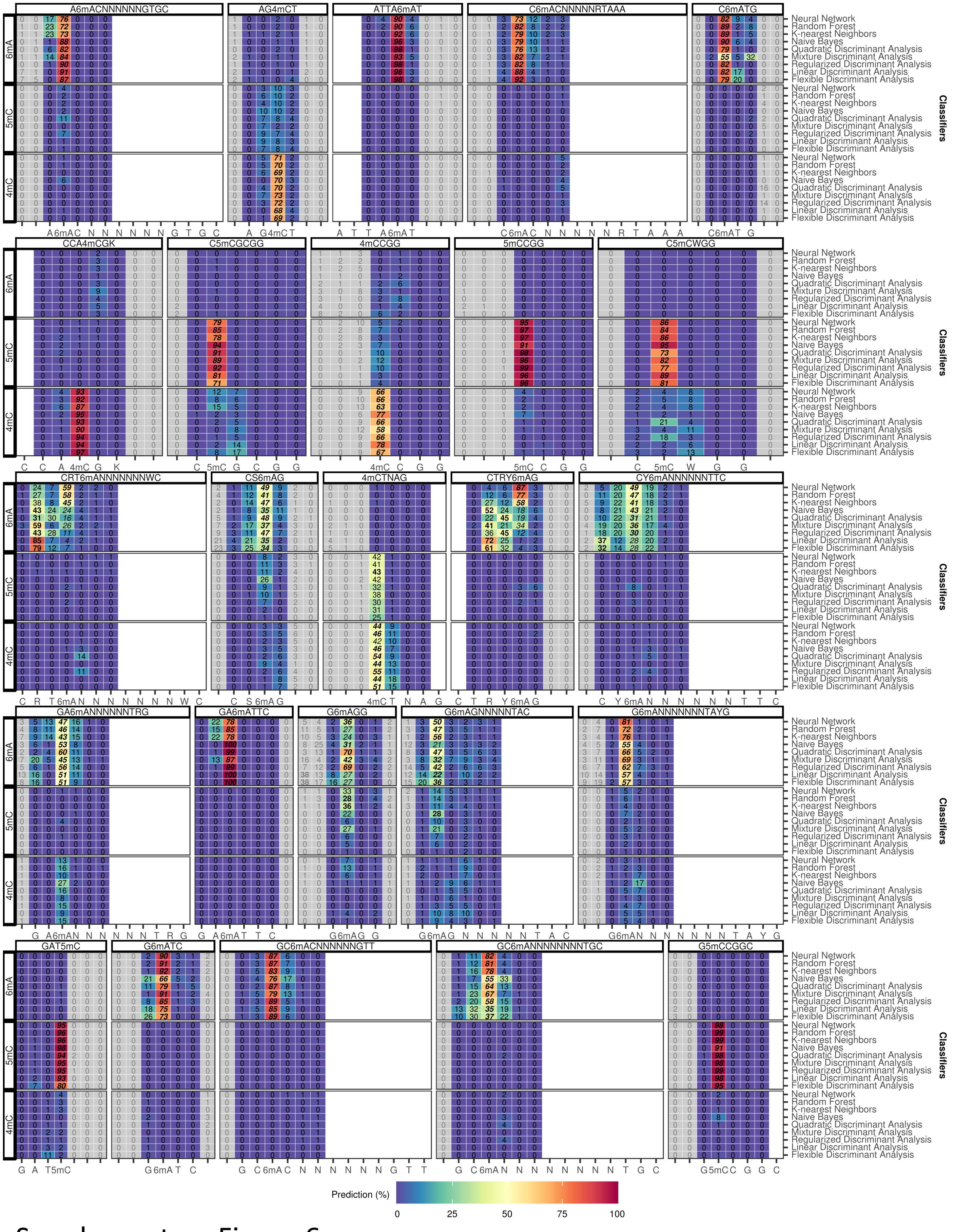


Supplementary Figure 6

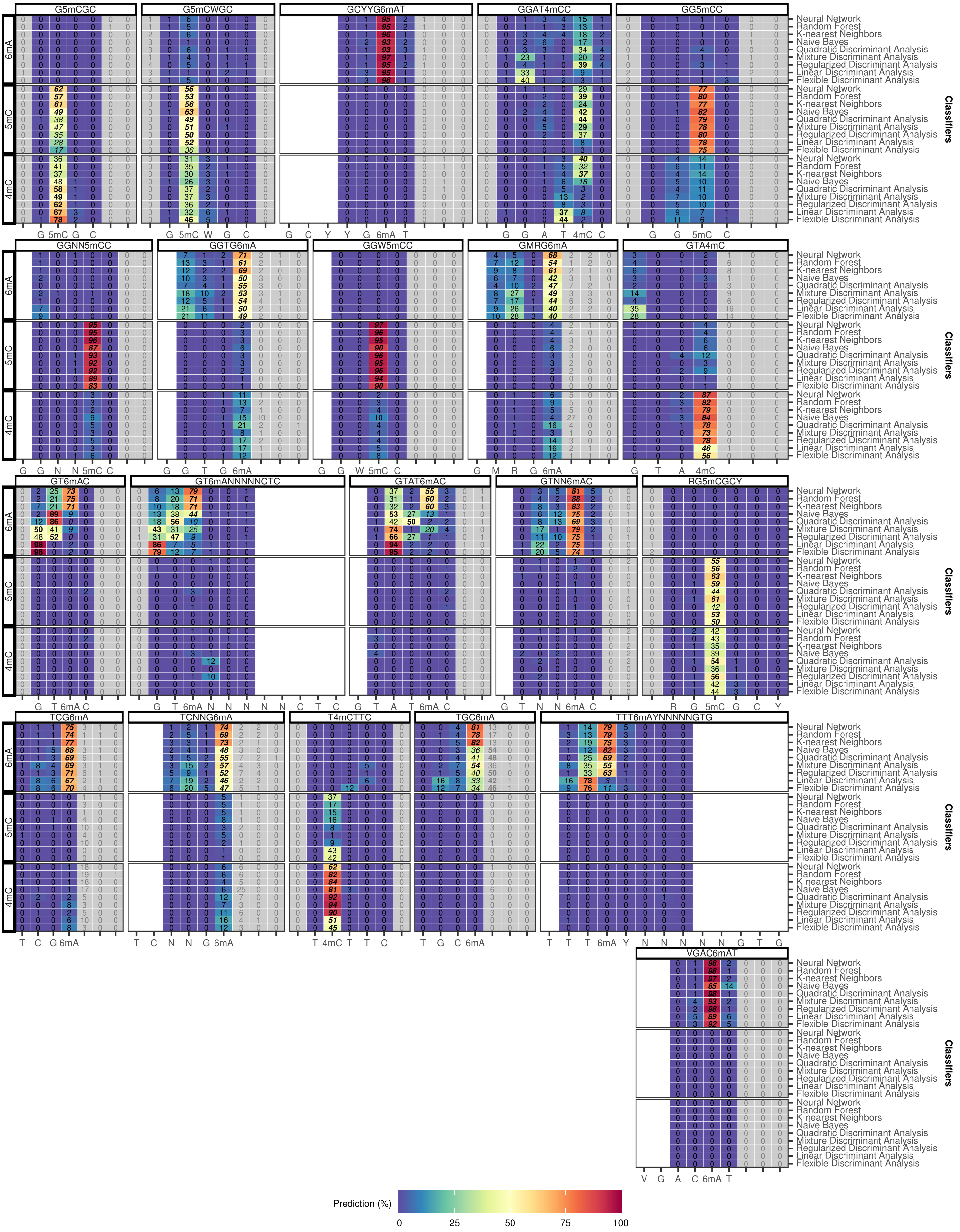


Supplementary Figure 7

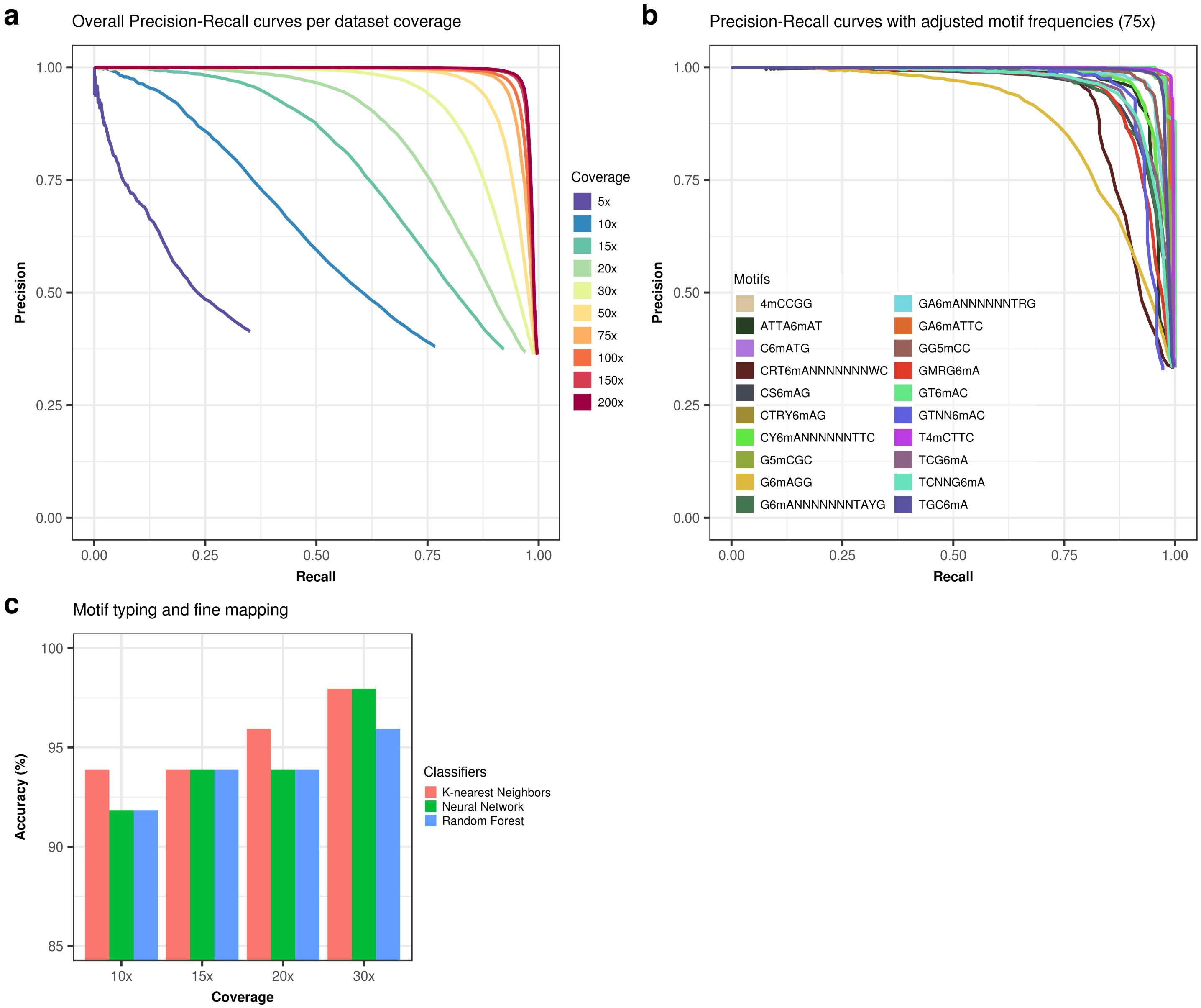


Supplementary Figure 8

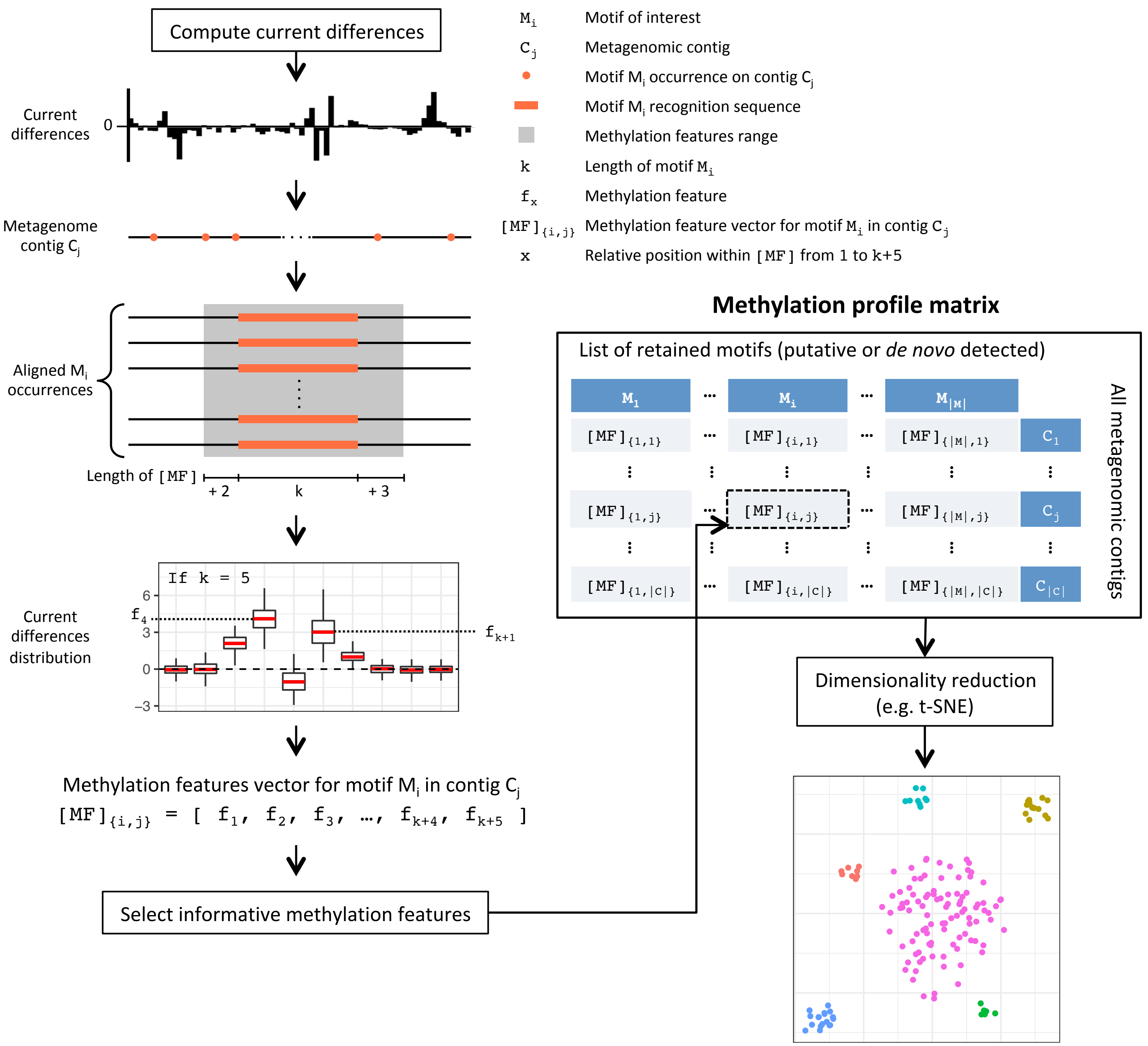


### Supplementary Figure 9

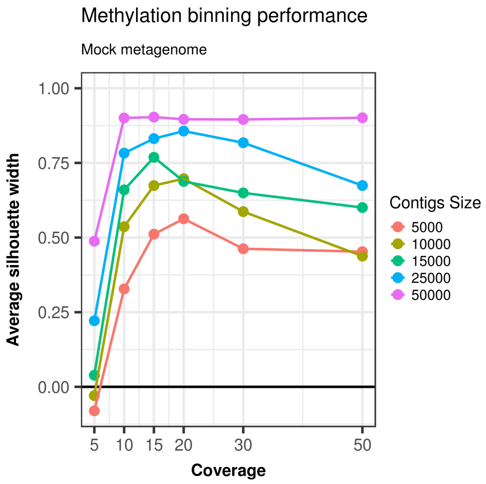


Supplementary Figure 10

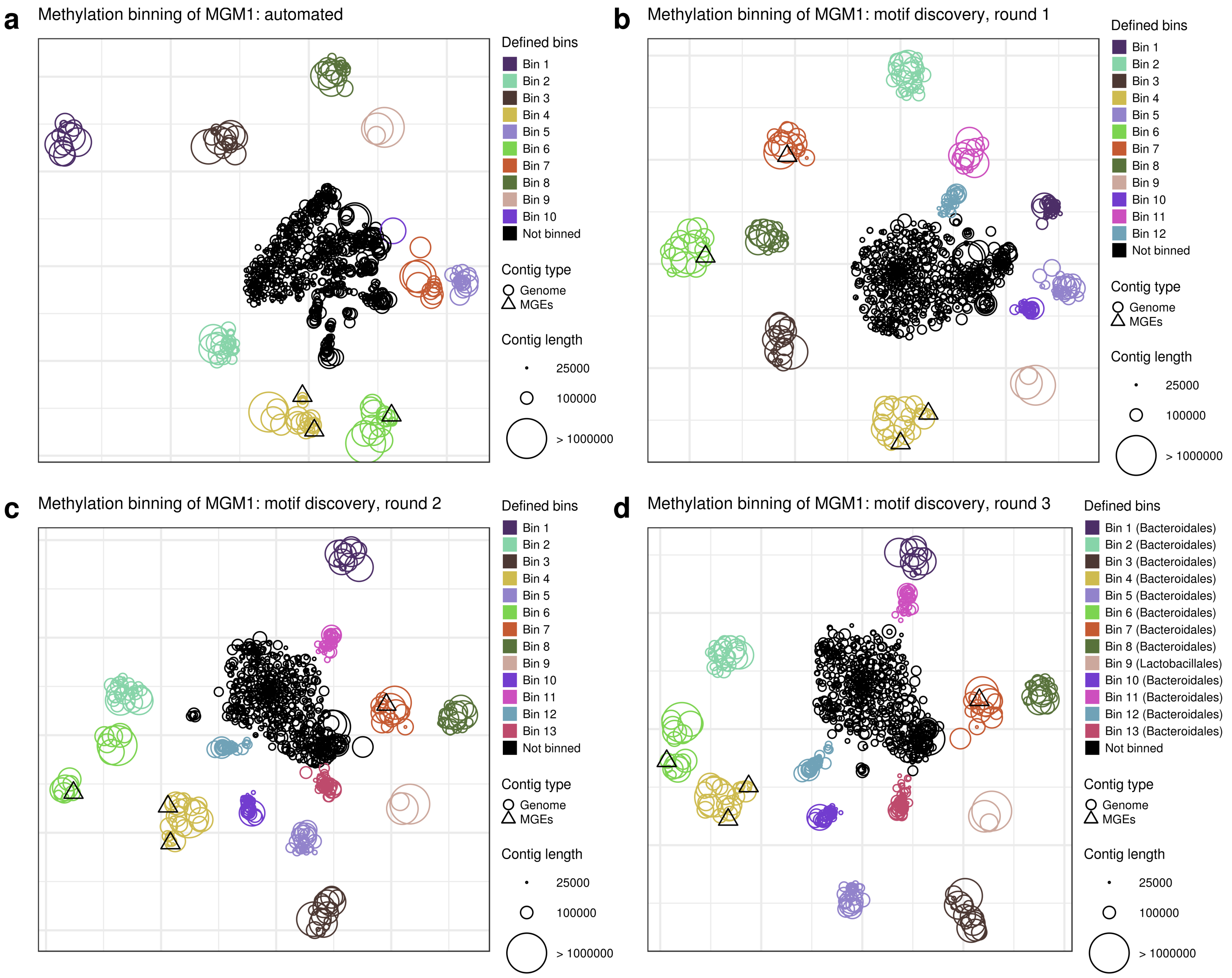


Supplementary Figure 11

**a** Methylation binning of MGM2: automated

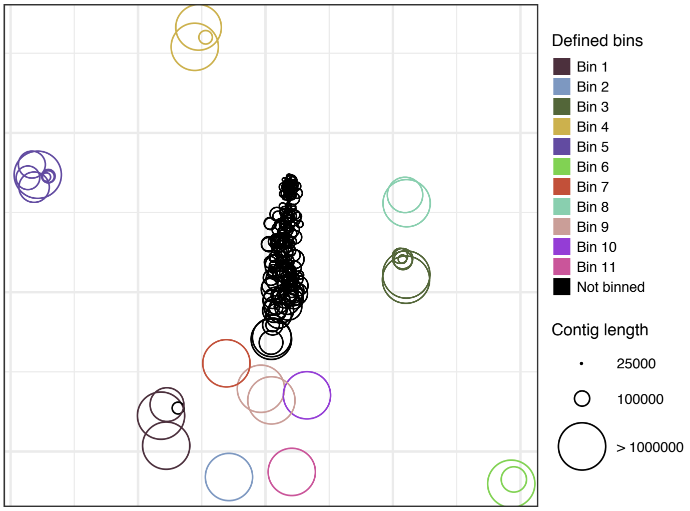


**b** Methylation binning of MGM2: motif discovery

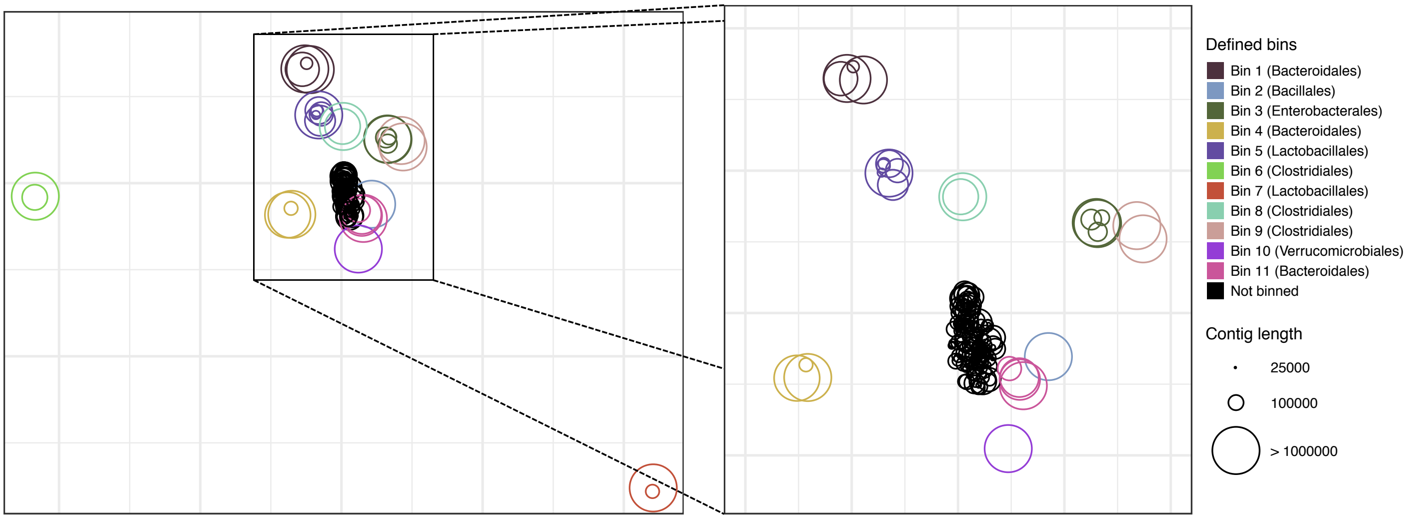


Supplementary Figure 12

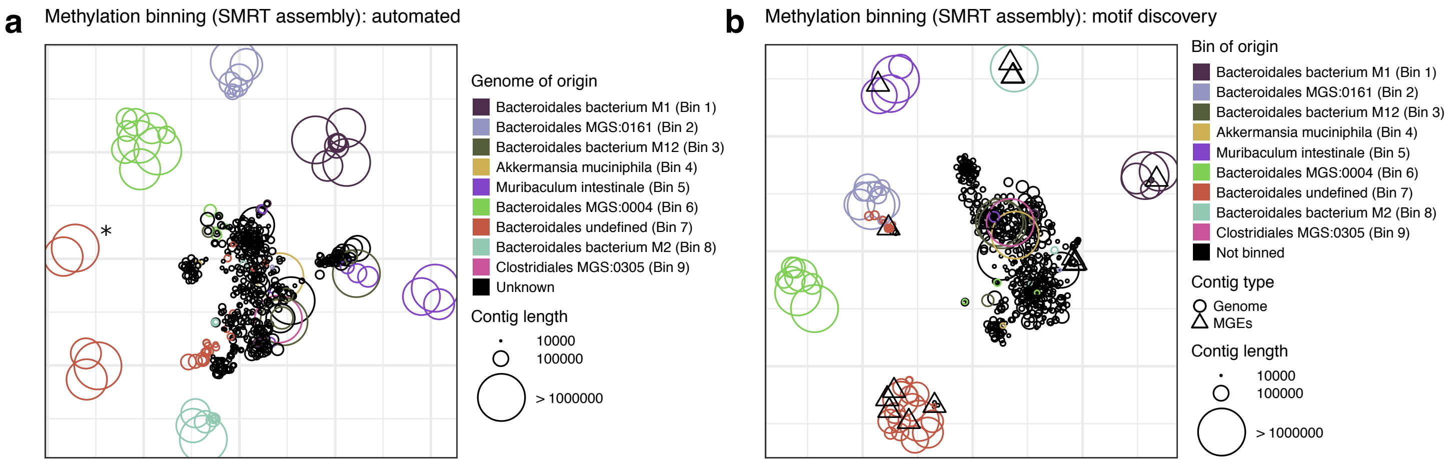


Supplementary Figure 13

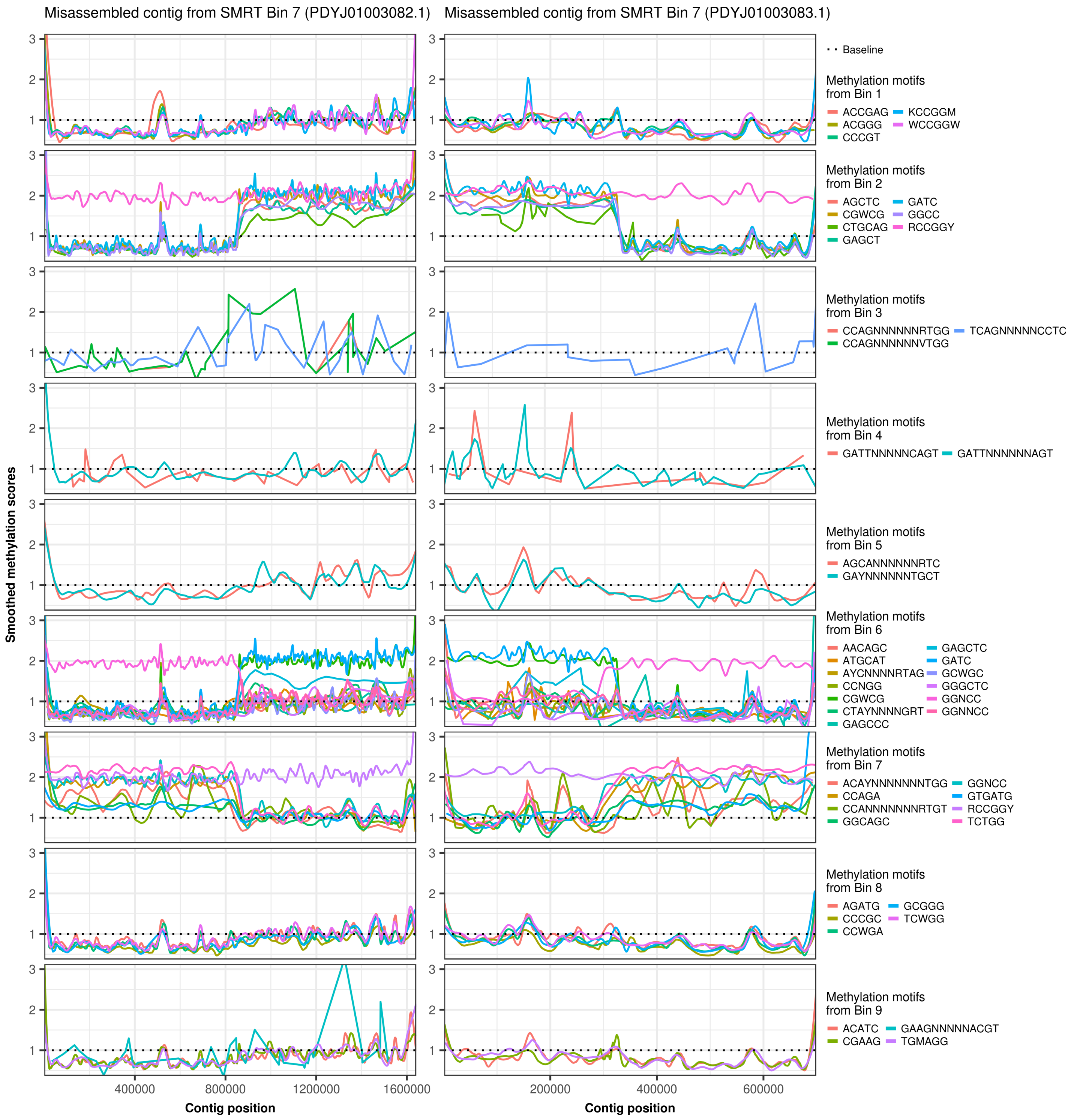


Supplementary Figure 14
